# CECR2 Drives Breast Cancer Metastasis by Suppressing Macrophage Inflammatory Responses

**DOI:** 10.1101/2020.09.10.291799

**Authors:** Meiling Zhang, Zongzhi Z. Liu, Keisuke Aoshima, Yangyi Zhang, Yongyan An, Asako Aoshima, Lok Hei Chan, Sabine M. Lang, Hongyi Sun, Zhenwei Tang, Sara J. Rutter, Carmen J. Booth, Veerle Bossuyt, Xiang Chen, Jon S. Morrow, Lajos Pusztai, David. L. Rimm, Mingzhu Yin, Qin Yan

## Abstract

Epigenetic and transcriptional changes are critical for metastasis, the major cause of cancer-related deaths. Metastatic tumor cells escape immune surveillance more efficiently than tumor cells in the primary sites, but the mechanisms controlling their immune evasion are poorly understood. We found that distal metastases are more immune inert with increased M2 macrophages compared to their matched primary tumors. Acetyl-lysine reader CECR2 is an epigenetic regulator upregulated in metastases and positively associated with M2 macrophages. CECR2 specifically promotes breast cancer metastasis in multiple mouse models, with more profound effect in the immunocompetent setting. Mechanistically, NF-κB family member RELA recruits CECR2 to activate *CSF1* and *CXCL1*, which are critical for macrophage-mediated immunosuppression at the metastatic sites. Furthermore, pharmacological inhibition of CECR2 bromodomain impedes NF-κB-mediated immune suppression by macrophages and inhibits breast cancer metastasis. These results reveal novel therapeutic strategies to treat metastatic breast cancer.

**Statement of Significance:** Comparison of matched primary breast tumors and distal metastases show that metastases are more immune inert with increased tumor promoting macrophages. Depletion or pharmacological inhibition of CECR2 inhibits breast cancer metastasis by suppressing macrophage inflammatory responses, nominating CECR2 as a promising therapeutic target for cancer metastasis.

## Introduction

Breast cancer is the most common cancer among women worldwide and the second leading cause of cancer-related deaths in the United States (1,2). Breast cancer is heterogeneous genetically and clinically, and genetic and epigenetic changes accumulate continuously during the clinical course of the disease (3). The major cause of cancer related deaths is breast cancer metastasis to distal organs, including lung, brain and bone (4–7). There are many treatment options for patients with metastatic breast cancer, but despite of recent advances in treatment metastatic breast cancer remains incurable (2). Thus, there is an urgent need to identify new drug targets for the development of effective therapies.

Cancer metastasis is a multistep process of dynamic interactions between tumor cells and host microenvironment. The major steps are local invasion, intravasation, circulation, extravasation, and colonization at distant metastasis sites (8,9). Tumor cells not only activate immune tolerogenic signaling pathways, but also modulate tumor microenvironment by recruiting immune cells, endothelial cells, and fibroblasts, which contribute to cancer progression and metastasis (10–13). We have recently shown that metastatic breast cancers have a more immunologically inert tumor microenvironment than primary tumors (14). Several studies have shown that enhancing immune infiltration and activation leads to better treatment outcomes, providing important evidence for the development of more effective breast cancer immunotherapies (15–18). Tumor-associated macrophages (TAMs) are a major cell population in the tumor microenvironment and play key roles in carcinogenesis (19). TAMs are induced by signals to polarize into either classically activated M1 macrophages with a pro-inflammatory role, or alternatively activated M2 macrophages that promote tumor growth and metastasis (20–22). Research on TAMs has mainly focused on their roles in primary tumors; more studies to investigate the roles of TAMs as promoters or inhibitors of the metastatic cascade are needed (23).

Cat eye syndrome chromosome region candidate 2 (CECR2) was identified as a candidate gene for Cat Eye Syndrome (24). CECR2 contains a DDT domain, BAZ domain and bromodomain, which can recognize acetyl lysine residues and function in chromatin remodeling by interacting with SNF2L and SNF2H (25,26). CECR2 was also shown to play critical roles in DNA damage responses (27), neurulation (25) and spermatogenesis (26). The bromodomain of CECR2 has been predicted to be highly druggable (28), and two highly potent and specific CECR2 inhibitors GNE-886 and NVS-CECR2-1 have respectively been developed by Genentech (29) and the Structural Genomics Consortium (SGC) with Novartis (http://www.thesgc.org/chemical-probes/NVS-1). However, the specific functions of CECR2 in cancer, especially in the context of cancer immunity and metastasis remain unclear and limit the applications of these inhibitors.

NF-κB is a protein complex and has five family members, RELA/p65, c-REL, RELB, NF-κB1 (p50), and NF-κB2 (p52). These transcription factors form homodimers or heterodimers to activate their target gene transcription (30,31). IκBα binds to these dimers and renders them transcriptionally inactive in the absence of stimuli. Multiple signals, including cytokines, growth factors, DNA damage, oncogenic stress, could activate NF-κB signaling pathway (30). The canonical NF-κB pathway can be activated by the IKK complex, which phosphorylates IκBα, leading to the detachment of IκBα from NF-κB, release of NF-κB dimers into the nucleus, and activation of target gene transcription (32,33). Many cofactors are involved in NF-κB transcriptional activation, including histone acetyltransferase (HAT) p300, CBP, SRC-1, and TIF2. These cofactors promote the formation of an initiation complex by linking NF-κB with the transcriptional machinery (34–36). NF-κB activates immune and inflammatory responses, as well as cellular adhesion, metabolism, cell survival and proliferation (37,38). The aberrant activity of NF-κB in tumors is normally associated with increased cell proliferation, suppressed apoptosis, enhanced angiogenesis, and increased metastasis.

Herein, we profiled the transcriptomes of 13 matched primary and metastatic breast tumors and analyzed the immunological differences by comparing immune escape genes and immune-oncology targets. We found that the ratio of M2 macrophages was increased in metastatic tumor microenvironment. CECR2 was identified as the top epigenetic regulator of this increase as its mRNA levels correlated with M2 macrophage ratios. CECR2 knockout significantly decreased metastasis in multiple mouse breast cancer models. RNA-seq analysis revealed that CECR2 was essential for activation of NF-κB signaling in metastatic breast cancer cells. Mechanistically, CECR2 formed a complex with RELA through its bromodomain on the promoters of NF-κB target genes including *CSF1* and *CXCL1* to induce their expression. Furthermore, CECR2 stimulated the recruitment and polarization of tumor associated macrophages through CSF1 secreted by cancer cells, creating an immunosuppressive tumor microenvironment. Pharmacological inhibition of CECR2 suppressed NF-κB target genes and M2 macrophage polarization, and inhibited breast cancer metastasis. Taken together, our work establishes CECR2 as a novel epigenetic regulator of breast cancer metastasis and nominates it as a promising therapeutic target for the treatment of metastatic breast cancer.

## Results

### Immunological differences between metastatic and primary breast tumors

The tumor microenvironment plays key roles in shaping cancer metastasis and in determining treatment responses (39). By analyzing 730 immune-related genes using Nanostring technology, we showed recently that metastatic breast cancers have a more immunologically inert tumor microenvironment than primary tumors (14). However, it is poorly understood how this tumor microenvironment is controlled. To characterize the immune microenvironment differences more extensively and to identify regulators of tumor immune microenvironment and drivers of metastasis, we compared transcriptomes of 13 pairs of matched primary and distant metastatic breast cancer tumor samples using RNA sequencing (RNA-seq) analysis (Figure 1A). The median age of these patients was 51 years, and their median overall survival time was 4 years (Supplemental Table 1). Six patients had ER positive tumors, while seven patients had ER negative tumors. Tumor metastases for these patients were found in different locations, including ovary, lung, brain, liver, spine, esophagus, skin, stomach, fallopian tubes and soft tissue. Hierarchical clustering analysis revealed that all tumors from ER positive patients were clustered into one group, while ER negative tumors clustered separately (Supplemental Figure 1A). These results also indicated that the gene expression profiles of primary and metastatic tumors from the same patient were clustered together, despite their divergent locations. We found 930 significantly differentially expressed genes, among which 627 genes were significantly downregulated and 303 genes were significantly upregulated in the distant metastases versus the primary tumors (Supplemental Table 2).

**Figure 1.**
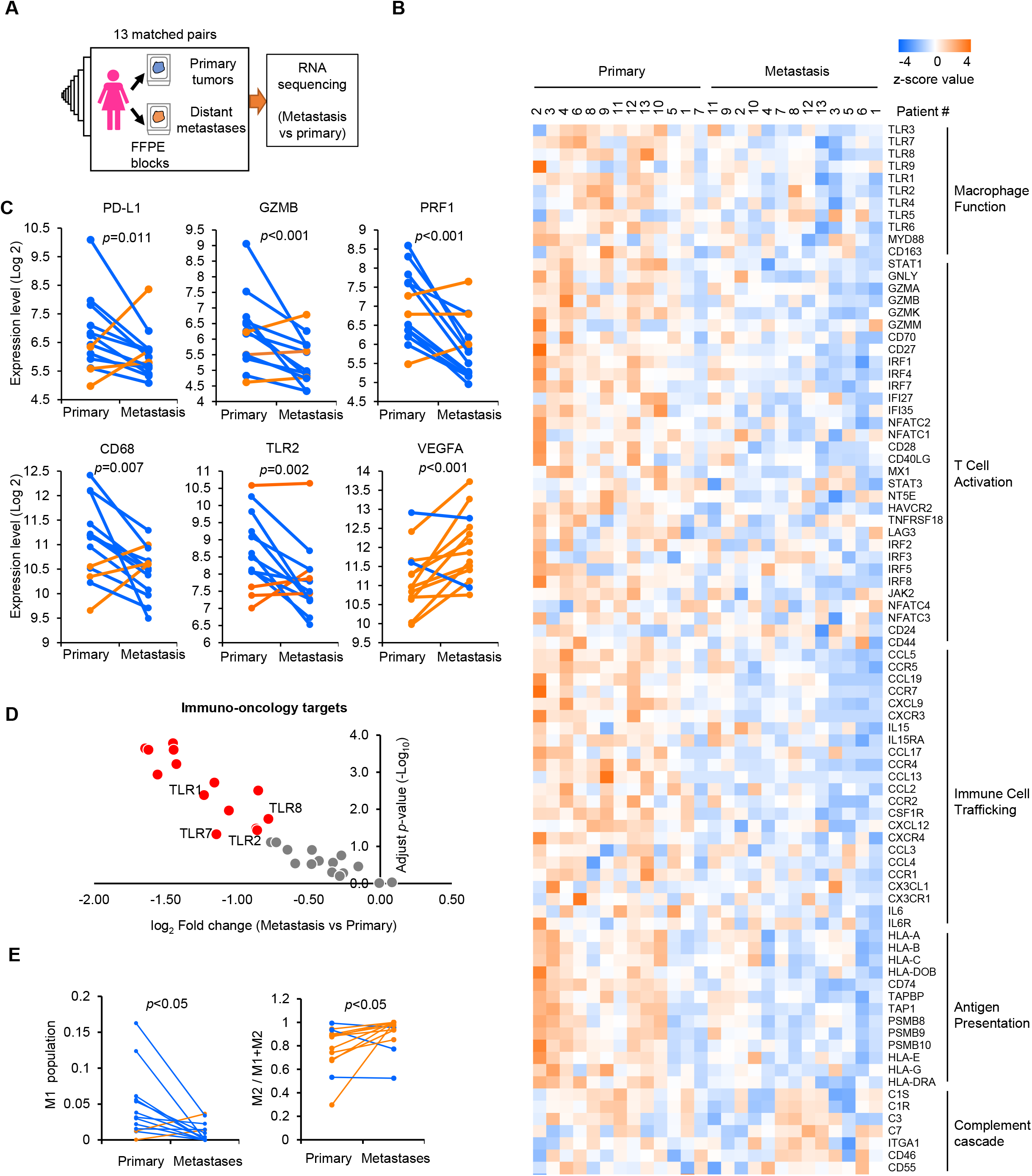
Immunological differences in metastatic and primary breast cancer. (**A**) Matched primary tumors and distal metastases from 13 breast cancer patients were collected and deregulated genes were analyzed by comparing distal metastases with matched primary tumors using RNA-sequencing (RNA-seq) analysis. (**B**) Heat map of the expression of representative immune genes of tolerance mechanisms in 13 pairs of primary and matched metastatic breast cancer tumor samples. (**C**) Tumor infiltrating lymphocyte marker and macrophage related gene expression in matched pairs of primary and metastatic breast tumor samples. Yellow lines marks the samples with increased expression in metastasis while blue lines marks the ones with decreased expression. (D) Volcano plot of downregulated immune-oncology targets in matched metastatic samples compared with primary breast tumors. (E) Population of M1 macrophages and ratio of M2 macrophages to total Macrophages in primary and matched metastatic breast cancer samples. Yellow lines marks the samples with increased number in metastasis while blue lines marks the ones with decreased number.

RNA-seq data showed that the majority of immune-related genes were downregulated in the metastatic tumors comparing to the matched primary tumors, especially the genes in macrophage function and T cell activation (Figure 1B). The anti-tumor immune response and activation markers, including PD-L1, Granzyme B (GZMB) and perforin (PRF1), all decreased in the metastasis tumor microenvironment (Figure 1C). Interestingly, genes associated with inflammatory macrophages, such as CD68 and TLR2, were downregulated, while VEGFA, contributing to cancer metastasis and M2 macrophage polarization, was upregulated in metastatic tumor microenvironment (Figure 1C). We also found 14 out of 29 immuno-oncology targets genes were significantly downregulated in metastatic tumors compared to their matched primary tumors, in which four genes (TLR1, TLR8, TLR2 and TLR7) are associated with macrophage functions (40,41), three genes (CCR4, CXCL12 and CXCR4) are associated with immune cell trafficking, and two genes (CTLA-4 and CD27) are involved in immune checkpoint function (Figure 1D, Supplemental Table 3). To understand the immune cell composition differences in matched primary and metastatic tumor microenvironment, we analyzed the RNA-seq data using CIBERSORTx (42). The major components of immune cells from CIBERSORTx analysis are macrophages, CD4 T cells and B cells in tumor microenvironment (Supplemental Table 4). Intriguingly, the M1 macrophage population is significantly decreased and the ratio of M2 macrophages to total macrophages increased in metastasis tumor (Figure 1E, Supplemental Figure 1B). However, the total macrophages showed no difference between primary tumors and matched metastases, as well as CD8+ and CD4+ T cells, NK cells, dendritic cells, and neutrophils (Supplemental Figure 1, C-H). These results indicate that the population variation of macrophages, especially the M2 ratio, is the major immunological difference between primary and metastasis breast cancer tumor microenvironment.

### CECR2 expression is associated with breast cancer metastasis

Epigenetic and transcriptional changes have been implicated in metastatic progression. We focused on our attention on epigenetic regulators that were altered in the metastatic niche. To this end, we compared the list of differentially expressed genes with the list of genes involved in epigenetic regulation that we compiled (Supplemental Table 5) by combining the epigenetic gene lists in the literature (43,44) and at the SGC website. Among the 24 significantly deregulated epigenetic genes with fold change more than 1.5 (Figure 2, A and B, and Supplemental Table 6), PPARGC1A (gene encoding PGC-1α) was reported to promote breast cancer metastasis (45) and it was also upregulated in our screening of breast cancer patients. Beyond this positive control, we found several additional potential novel epigenetic or transcriptional regulators of breast cancer metastasis, including CECR2, FOXP family proteins, nuclear body proteins, DNA methylation regulators, and PR-domain proteins.

**Figure 2.**
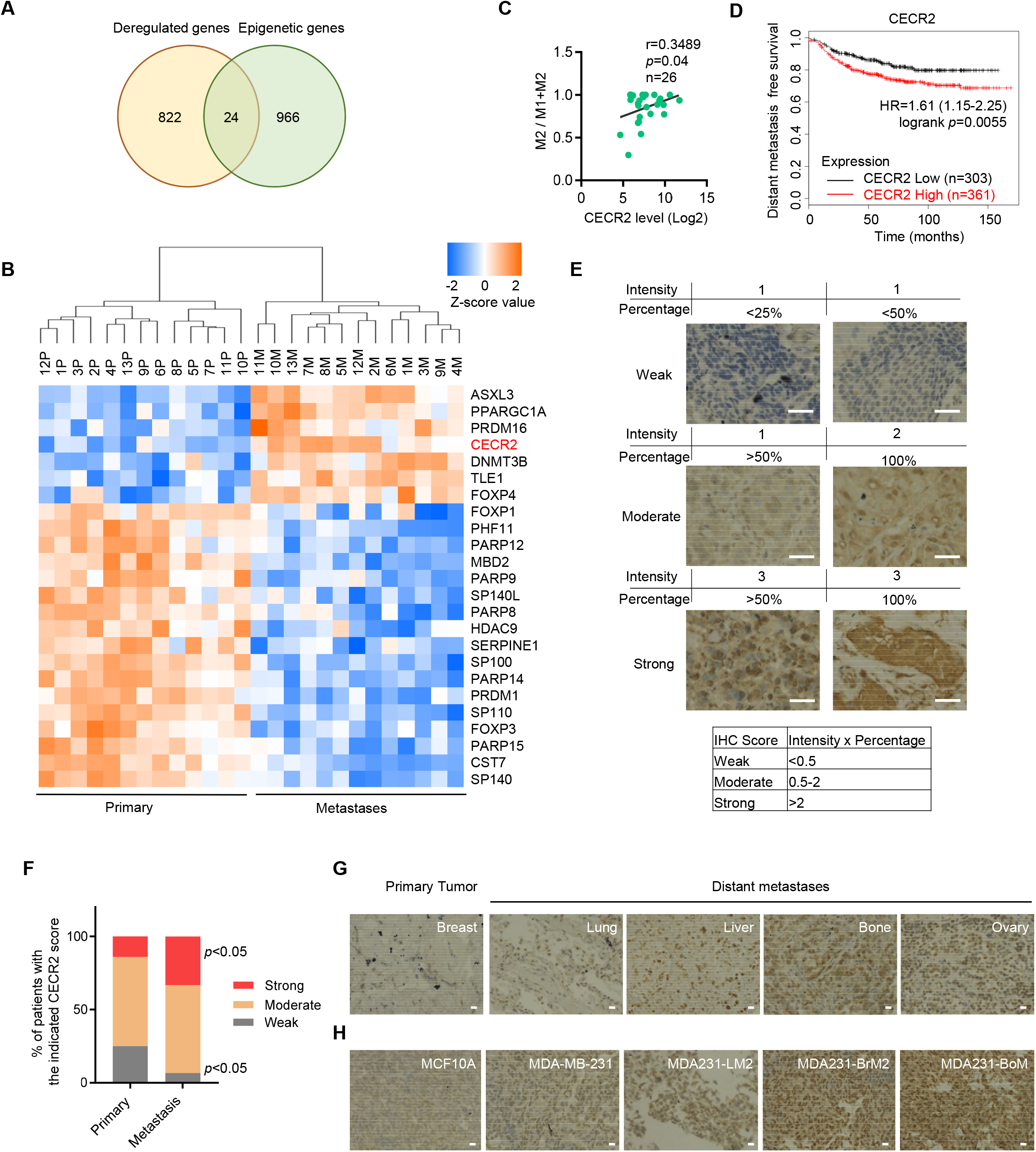
CECR2 is correlated with M2 macrophages and is highly expressed in breast cancer metastases. (**A**) Venn diagram showing deregulated epigenetic genes with significantly changed mRNA expression level (fold change >1.5 and <−1.5) by RNA-seq in metastatic samples compared to primary samples. (**B**) Heat map of the significantly deregulated epigenetic genes. CECR2 is highlighted in red. (**C**) RNA-seq data of matched primary tumor and distal metastases from 13 breast cancer patients were analyzed by CIBERSORTx and immune cell composition of complex tissues were characterized from their gene expression profiles. The correlation of M2 ratios with CECR2 expression levels was shown. Spearman correlation coefficient and One-tailed probability *p* value were calculated. (**D**) Kaplan-Meier (KM) plotter analysis showing association of CECR2 mRNA levels with distant metastasis free survival of breast cancer patients using the best cutoff. Hazard ratio (HR) and log-rank *p* values were calculated. (**E**) CECR2 Immunohistochemistry (IHC) staining of tumor tissue microarray with 59 pairs of matched primary and metastatic breast cancer samples. Representative figures were shown. Scale bars: 100 μm. (**F**) CECR2 IHC scores were quantified by multiplying the intensity of the signal and the percentage of positive cells. The IHC staining of tumors were scored as weak (score< 0.5), moderate (score between 0.5 and 2) and strong (score >2). Percentage of patient samples with strong CECR2 level in metastatic tumors vs that in primary tumor, *p*<0.05. Percentage of samples with weak CECR2 level in metastatic tumors vs that in primary tumor, *p*<0.05. (**G**) CECR2 IHC staining of matched primary and multiple distant metastasis samples from a single breast cancer patient. Scale bars: 100 μm. (**H**) CECR2 IHC staining of MCF10A, MDA-MB-231 and its metastatic derivatives (MDA231-LM2, MDA231-BrM2 and MDA231-BoM). Scale bars: 100 μm.

The analysis of transcriptome expression in primary and metastasis breast cancer tumor indicates that metastatic tumor microenvironments are more inert in breast cancer (Figure 1). To investigate how epigenetic change regulates immune microenvironment during breast cancer metastasis, we analyzed the correlation of M2 macrophage ratio with the expression of each dysregulated epigenetic factor. The expression of 11 epigenetic factors significantly correlated with the ratio of M2 macrophage, among which CECR2 is the only gene that was overexpressed and showed positive correlation with the ratio of M2 macrophage (Figure 2C, Supplemental Table 7). Consistent with these results, Kaplan-Meier plotter analysis (46) showed that high CECR2 mRNA levels were associated with poor distant metastasis free survival of breast cancer patients overall and in ER^+^ and HER2^+^ breast cancer subtypes (Figure 2D, and Supplemental Figure 2A). Similar results were found in gastric and ovarian cancer cohorts (Supplemental Figure2, B and C).

Herein, we have focused on CECR2 as it is a novel targetable epigenetic factor for breast cancer metastasis. Increased CECR2 mRNA levels in distant metastases were confirmed by RT-qPCR assays (Supplemental Figure 2D). We further examined CECR2 protein expression by IHC staining of a tissue microarray comprised of 59 pairs of matched human primary tumors and distant metastases (Supplemental Table 8, expanded from previously described (14). Two pathologists independently evaluated CECR2 expression levels by the IHC scores (stain intensity score multiplied by the percentage of positive tumor cells) and found that higher CECR2 protein levels were more frequently observed in cancer cells in the distant metastases than in the primary tumors (Figure 2, E and F, and Supplemental Table 8). To characterize the relationship of CECR2 expression with the location of metastases, we performed IHC staining with breast cancer samples taken from one patient with multiple metastatic sites, including lung, liver, bone and ovary. We found that all the metastatic samples have higher levels of CECR2 expression, with the highest levels in the bone and ovary (Figure 2G). We also compared immortalized MCF10A breast epithelial cells, triple negative MDA-MB-231 breast cancer cells (MDA231) and MDA231-derived metastatic cell lines, including MDA231-LM2 (LM2), MDA231-BrM2 (BrM2) and MDA231-BoM (BoM) cells. These three MDA231 metastatic cell lines were derived by *in vivo* selection, with increased metastatic activity to the lungs, brain and bones, respectively, compared with their parental cells (47–49). CECR2 protein was expressed at a higher level in MDA231 cells than in MCF10A cells (Figure 2H). All three MDA231 derivatives have increased CECR2 protein levels compared with the parental MDA231 cells (Figure 2H). Taken together, CECR2 level is correlated with increased metastatic potential.

### CECR2 is critical for migration, invasion and metastasis

To dissect the roles of CECR2 in metastasis, we first generated polyclonal LM2 cell lines with stable CECR2 knockout (CECR2 sg) or non-targeting control (Control) using clustered regular interspaced short palindromic repeats (CRISPR)/CRISPR-associated protein 9 (Cas9) system (50) (Figure 3A). The firefly luciferase was engineered into these LM2 cells to monitor the metastasis signal *in vivo* by a live imaging system (48). Depletion of CECR2 has no effect on cell proliferation in both WST1 cell proliferation and colony formation assays (Supplemental Figure 3, A and B). Migration and invasion through tissue basement membrane is one of the key steps of metastasis. We examined the effects of CECR2 depletion on migration and invasion of LM2 cells using scratch assay, transwell migration and invasion assays. We found that CECR2 depletion dramatically decreased migration and invasion capability of LM2 cells (Figure 3, B and C, and Supplemental Figure 3C).

**Figure 3.**
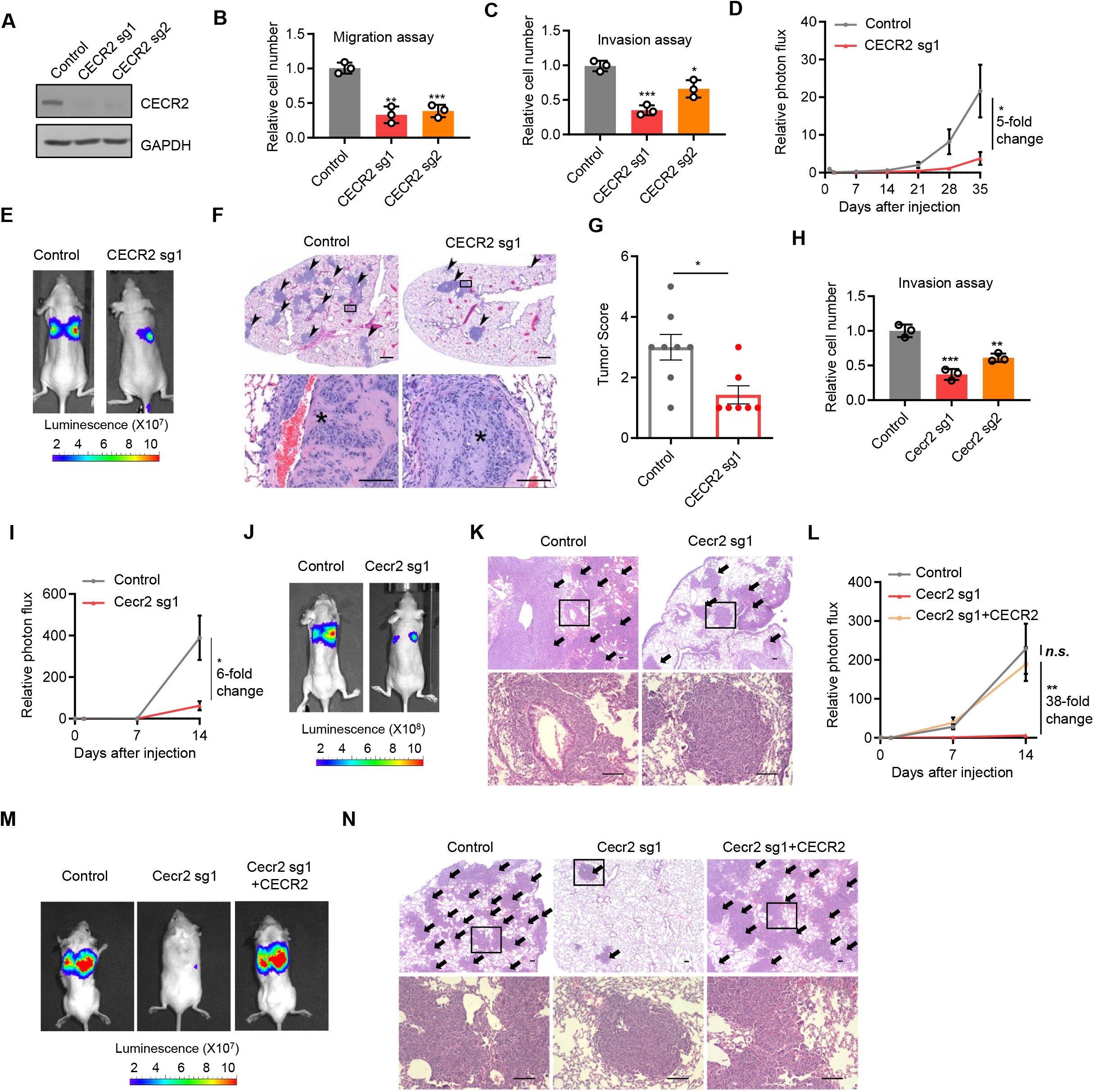
CECR2 is required for invasion and metastasis. (**A**) Western blot analysis of control and CECR2 knockout (sg1 and sg2) LM2 cells. (**B**and **C**) Transwell migration (**B**) and invasion (**C**) assays comparing control and CECR2 knockout LM2 cells. (**D**) Normalized bioluminescence signals of lung metastases in nude mice with tail vein injection of control (n=8) or CECR2 knockout LM2 cells (n=7). (**E**) Representative bioluminescence images of mice in (**D**) at week 5. (**F**) H&E staining of the lungs from mice in (**D**) at week 5. Scale bars: 500 μm for the upper panel and 100 μm for the lower panel. (**G**) Tumors were scored based on the percentage of tumors in the lungs with the parameter as **Figure S3E**. (**H**) Transwell invasion assays comparing control and Cecr2 knockout 4T1 cells. (**I**) Normalized bioluminescence signals of lung metastases in immunodeficient BALB/c nude mice with tail vein injection of control (n=6) and Cecr2 knockout 4T1 cells (n=7). (**J**) Representative bioluminescence images of mice in (**I**) at week 2. (**K**) H&E staining of the lungs from mice in (**I**) at week 2. Scale bars: 200 μm. (**L**) Normalized bioluminescence signals of lung metastases in immunocompetent BALB/c mice with tail vein injection of control 4T1 (n=10), cecr2 knockout 4T1 (n=10) and cecr2 knockout 4T1 with CECR2 reconstituted expression (n=10). (**M**) Representative bioluminescence images of mice in (**L**) at week 2. (**N**) H&E staining of the lungs from mice in (**L**) at week 2. Scale bars: 200 μm. \**p* < 0.05, ** *p* < 0.01, *** *p* < 0.001, *n.s.*, not significant. Representative data from triplicate experiments are shown, and error bars represent SEM.

To determine the roles of CECR2 in metastasis *in vivo*, LM2 cells with stable CECR2 knockout or control were injected into athymic nude mice through tail vein. We found that CECR2 knockout led to about 5-fold decrease in lung colonization capability of LM2 cells and extended survival of tumor bearing mice using bioluminescence signal as the end point (Figure 3, D and E, and Supplemental Figure 3D). Consistently, histological analysis of mouse lungs showed that CECR2 knockout LM2 cells formed fewer tumor lesions than control cells (Figure 3F). Quantification of these lesions showed that CECR2 knockout strongly decreased tumor score in the lungs (Figure 3G, and Supplemental Figure 3E).

We next extended our studies using 4T1 mouse triple negative breast cancer cell line with stable Cecr2 knockout and stable expression of firefly luciferase (Supplemental Figure 4, A and B). Consistent with the results in LM2 cells, Cecr2 depletion decreased cell invasion, but not tumor cell proliferation (Figure 3H, and Supplemental Figure 4, C-E). Cecr2 depletion in 4T1 cells suppressed their metastatic potential to the lungs by about 6-fold and extended the survival of tumor bearing BALB/c nude mice using bioluminescence signal as the end point (Figure 3, I and J, and Supplemental Figure 4F) and histological analysis (Figure 3K).

We found that metastatic sites have different tumor immune microenvironments from the primary tumors (Figure 1) (14), thus we examined the effects of Cecr2 loss in an immunocompetent setting. To eliminate the off-target effect of Cecr2 sgRNA, we also restored CECR2 expression in Cecr2 knockout 4T1 cells using human CECR2 (Supplemental Figure 4G). We then injected these cells into BALB/c mice through tail vein and monitored their ability to colonize the lungs. Cecr2 knockout led to about 38-fold decrease of lung metastasis and significantly extended the survival of tumor bearing mice using bioluminescence signal as the end point, and restored expression of CECR2 completely rescued the phenotype (Figure 3, L-N, and Supplemental Figure 4, H and I). Of note, suppression of metastasis by Cecr2 loss in immunocompetent mice (38-fold) is more profound than that in immunodeficient mice (6-fold), suggesting tumor immune microenvironment contributes significantly to this difference. Consistent with the role of CECR2 in distal metastasis, Cecr2 depletion in 4T1 cells did not affect their tumor growth rate in mammary fat pads of immunocompetent mice, but significantly decreased spontaneous lung metastasis (Supplemental Figure 4, J and K).

### Regulation of the NF-κB pathway by CECR2

To investigate the underlying molecular mechanisms by which CECR2 modulates breast cancer metastasis, we examined the transcriptome changes in LM2 cells after CECR2 knockout using RNA-seq analysis. We observed 1,051 significantly upregulated and 1,440 significantly downregulated genes in LM2 cells with CECR2 sg1 (Supplemental Table 9). Similarly, there were 1,708 significantly upregulated and 1,772 significantly downregulated genes in LM2 cells with CECR2 sg2 (Supplemental Table 10). By gene set enrichment analysis (GSEA), we found 8 shared down-regulated hallmark pathways and 2 shared upregulated hallmark pathways by CECR2 sg1 and sg2 (Figure 4, A and B, and Supplemental Figure 5, A and B, and Supplemental Table 11-14). The downregulated pathways include TNFA signaling Via NF-κB, inflammatory response, KRAS signaling, estrogen response and EMT pathways (Figure 4B, and Supplemental Figure 5, C-F). Most NF-κB response genes were suppressed by CECR2 knockout, including genes encoding cytokines CSF1, CSF2 and CXCL1 (Supplemental Figure 5E). The regulation of these NF-κB response genes by CECR2 was confirmed by RT-qPCR and western blot analysis of LM2 (Figure 4C, and Supplemental Figure 6A) and 4T1 cells (Figure 4D, and Supplemental Figure 6B).

**Figure 4.**
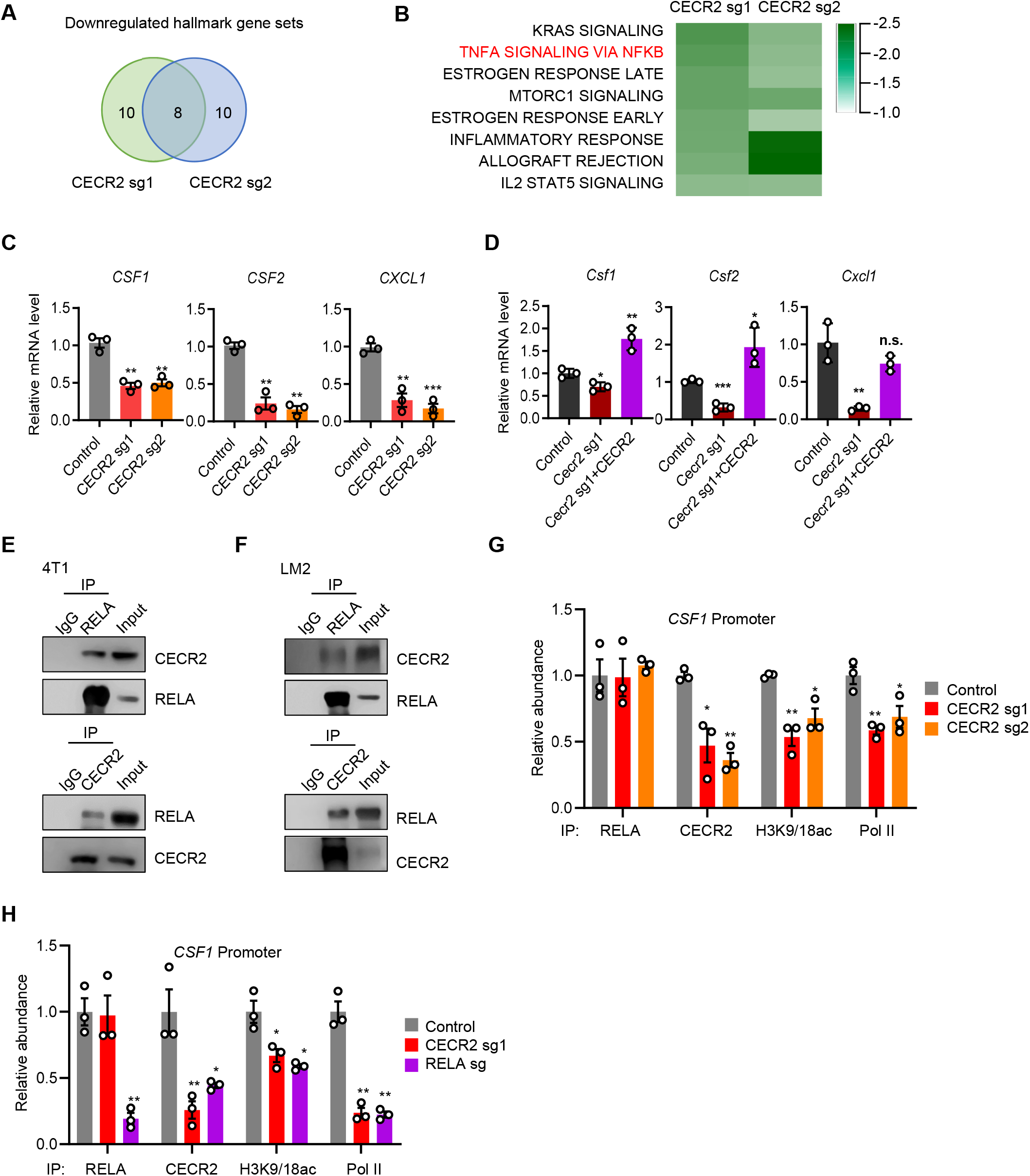
CECR2 interacts with acetylated RELA using its bromodomain to activate NF-κB response genes. (**A-B**) Gene set enrichment analysis comparing transcriptomes of CECR2 knockout (CECR2 sg1 and CECR2 sg2) with control LM2 cells. Venn diagram (**A**) showing the number of shared downregulated hallmark pathways (**B**). (**C**) RT-qPCR analysis of *CSF1, CSF2* and *CXCL1* in control and CECR2 knockout LM2 cells treated with 20 ng/ml TNF-α for 3 hours. (**D**) RT-qPCR analysis of *Csf1, Csf2* and *Cxcl1* in control 4T1, cecr2 knockout 4T1 and cecr2 knockout 4T1 with CECR2 reconstituted expression after treatment with 20 ng/ml TNF-α for 3 hours. (**E-F**) Western blot analysis of cell lysates (input) and immunoprecipitates (IP) from 4T1 (**E**) and LM2 (**F**) cells stimulated with 20 ng/ml TNF-α for 0.5 hour with the indicated antibodies. (**G-H**) ChIP-qPCR analyses with the indicated antibodies of the *CSF1* promoter in control, CECR2 knockout (CECR2 sg1 and CECR2 sg2) (**G**), and RELA knockout (RELA sg) (**H**) LM2 cells stimulated with 20 ng/ml TNF-α for 0.5 hour. * *p* < 0.05, ** *p* < 0.01, *** *p* < 0.001. Representative data from triplicate experiments are shown, and error bars represent SEM.

### CECR2 binds to acetylated RELA to activate the NF-κB response genes

We then asked whether CECR2 loss affects the transcription factors that control the expression of NF-κB targeted genes. CECR2 knockout did not change the protein levels of NF-κB family members, including RELA/p65, p50, RELB, p52 and cREL in the cytosol and nucleus (Supplemental Figure 6C). Co-immunoprecipitation experiments showed that CECR2 interacts with RELA in both 4T1 and LM2 breast cancer cells endogenously (Figure 4, E and F) and in 293T cells exogenously (Supplemental Figure 6D). To determine the roles of the CECR2-RELA interaction on transcription of NF-κB targeted genes, we performed ChIP-qPCR analyses of CECR2, RELA, transcriptional activation mark (H3K9-18ac) and RNA Pol II at the promoters of NF-κB target genes CSF1 and CXCL1. Depletion of CECR2 or RELA significantly decreased the levels of H3K9-18ac and Pol II at the promoters of CSF1 and CXCL1 in both LM2 (Figure 4, G and H, and Supplemental Figure 6, E and F) and 4T1 cells (Supplemental Figure 6, G and H). CECR2 deletion has no effect on RELA binding to these promoters (Figure 4, G and H, and Supplemental Figure 6, E-H). In contrast, RELA depletion inhibited CECR2 binding (Figure 4H), suggesting that RELA recruits CECR2 to activate gene expression.

As CECR2 is a bromodomain containing protein and bromodomains interact with acetylated proteins, we asked whether CECR2 interacts with RELA by recognizing acetylated residues in RELA. Interestingly, it was shown that BRD4 bromodomain recognizes lysine-310 acetylation of RELA (51). Thus, we mutated lysine-310 of RELA and found that this mutation also dramatically decreased its interaction with CECR2 (Figure 5A). Deletion the bromodomain of CECR2 inhibited its interaction with RELA (Figure 5B). These results suggest that CECR2 interacts with acetylated RELA through its bromodomain. Consistently, CECR2 bromodomain specific inhibitors NVS-CECR2-1 and GNE-886 (29) blocked the interaction of CECR2 and RELA (Figure 5C). Both NVS-CECR2-1 and GNE-886 also reduced the expression of *CSF1/2* and *CXCL1* in a dose-dependent manner in metastatic breast cancer, lung cancer and melanoma cells (Figure 5, D and E, and Supplemental Figure 7, A and B), and impaired the migration and invasion capability of LM2 breast cancer cells (Figure 5, F and G, and Supplemental Figure 7, C and D). These results indicate that CECR2 bromodomain is crucial for acetylated RELA to activate their target genes in multiple cancers, and pharmacological targeting CECR2 bromodomain inhibits breast cancer migration and invasion.

**Figure 5.**
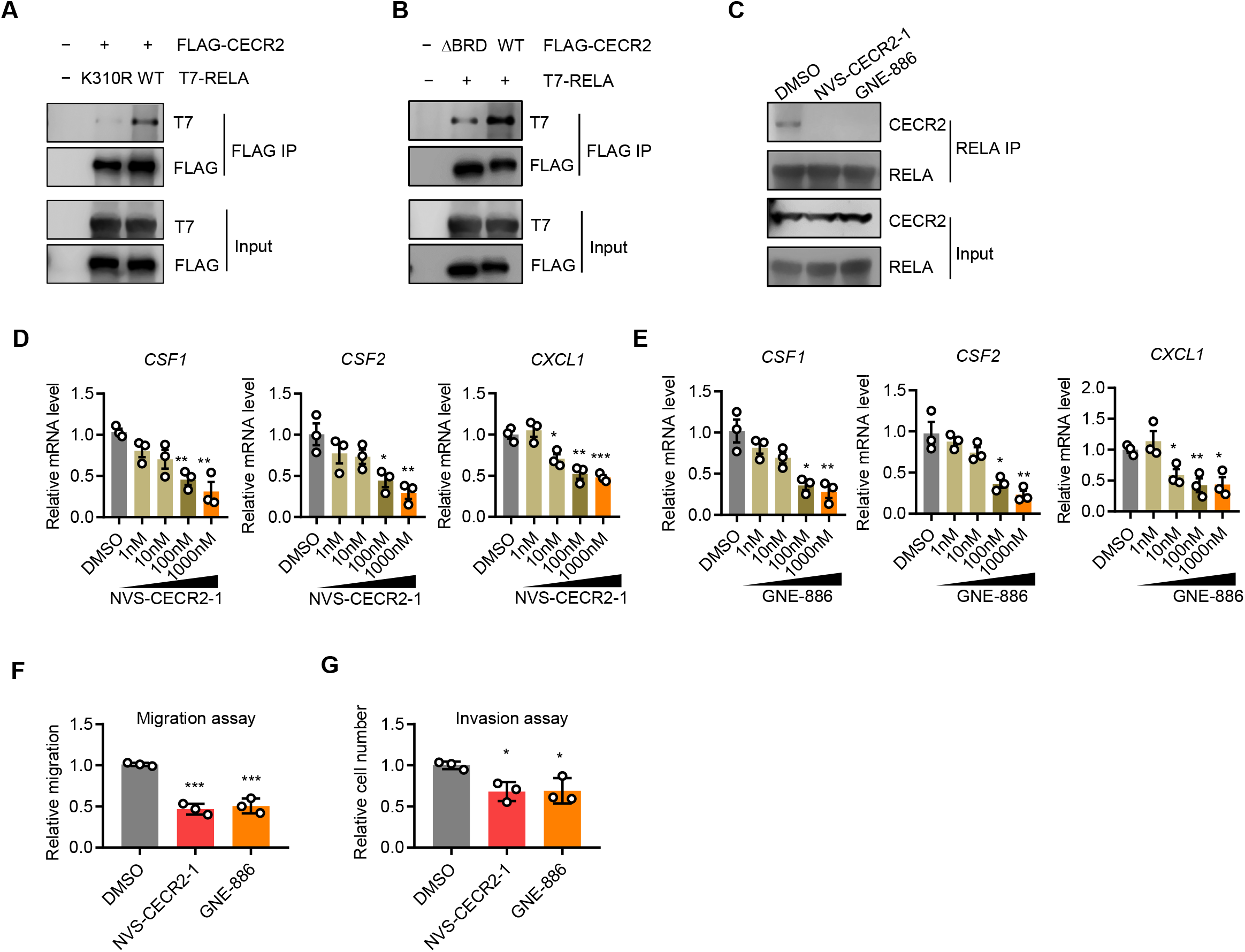
Pharmacological inhibition of CECR2 suppresses NF-κB response genes. (**A**) Western blot analysis of cell lysates (Input) and anti-FLAG immunoprecipitates (IP) from HEK293T cells transfected with the indicated combination of vectors expressing FLAG-CECR2, K310R mutated RELA and WT RELA. (**B**) Western blot analysis of cell lysates (Input) and anti-FLAG immunoprecipitates (IP) from HEK293T cells transfected with the indicated combination of vectors expressing WT FLAG-CECR2, FLAG-CECR2 mutant with bromodomain deletion (ΔBRD) and T7-RELA. (**C**) Western blot analysis of cell lysates (input) and anti-RELA immunoprecipitates (IP) from LM2 cells pretreated with control DMSO, CECR2 inhibitor 1 μM NVS-CECR2-1 or 1 μM GNE-886 for 2 days, and then stimulated with 20 ng/ml TNF-α for 0.5 hour. (**D-E**) RT-qPCR analyses of *CSF1, CSF2* and *CXCL1* in LM2 cells pretreated with the indicated concentration of NVS-CECR2-1 (**D**) or GNE-886 (**E**) for 2 days and then stimulated with 20 ng/ml TNF-α for 3 hours. (**F)**Scratch migration assays comparing the closure of wound healing distance in LM2 cells treated with DMSO, 1 μM NVS-CECR2-1 or 1 μM GNE-886 for 2 days. (**G**) Transwell invasion assays comparing LM2 cells treated with DMSO, 1 μM NVS-CECR2-1 or 1 μM GNE-886 for 2 days. * *p* < 0.05, ** *p* < 0.01, *** *p* < 0.001. Representative data from triplicate experiments are shown, and error bars represent SEM.

### CECR2 increases M2 macrophages in tumor immune microenvironment to drive tumor metastasis

We showed that M2 macrophage ratios are increased in metastatic tumors and are correlated with CECR2 levels (Figure 1E and Figure 2C). Moreover, CECR2 depletion decreased the expression of cytokines and chemokines, such as CSF1, CSF2 and CXCL1 (Figure 4, C and D, and Supplemental Figure 6, A and B). These cytokines/chemokines are involved in the monocytes/macrophages proliferation and differentiation in tumor microenvironment (52) and breast cancer metastasis (53). Therefore, we investigated whether CECR2 controls metastasis by regulating proliferation or polarization of tumor-associated macrophages. To examine the roles of tumor-intrinsic CECR2 on macrophage proliferation, we treated macrophages with the conditioned media (CM) from control and Cecr2 knockout 4T1 cells. The CCK8 cell proliferation assays showed that CM from control cells significantly promoted macrophage proliferation while CM from Cecr2 knockout cells abrogated the induction of macrophage proliferation (Figure 6A). We then studied the impact of tumor-intrinsic CECR2 on macrophage migration in a Boyden chamber co-culture system, in which tumor cells with or without Cecr2 depletion were placed into the lower chamber and macrophages were seeded into the upper chamber (Figure 6B). We found that Cecr2 knockout significantly decreased macrophage migration (Figure 6B). We next asked if tumor-intrinsic CECR2 affects macrophage polarization by treating macrophages with CM. We found that control CM strongly induced expression of M2 macrophage markers, while Cecr2 knockout CM are defective at inducing their expression (Figure 6C). To determine whether pharmacologically targeting CECR2 is a potential therapeutic option for metastatic breast cancer, we treated 4T1 tumor cells with different dosages of CECR2 bromodomain inhibitor NVS-CECR2-1 or GNE-886, then treated macrophages with CM from control and CECR2 inhibitor treated 4T1 cells. We found that the expression of M2 macrophage markers was suppressed by CM from CECR2 inhibitors treated cells in a dose dependent manner (Figure 6D). In contrast, treatment with NVS-CECR2-1 or GNE-886 on macrophage directly did not affect the expression of M2 macrophage markers (Supplemental Figure 8). To examine the roles of CECR2 in 4T1 tumor cells on macrophage polarization *in vivo*, we first performed flow cytometry analysis of the lung metastases from BALB/c nude mice implanted with 4T1 cells through tail vein. We showed that CECR2 loss in 4T1 cells decreased the number of macrophages and the ratio of M2 macrophages, but had minimal effect on the ratio of M1 macrophages and NK cells (Figure 6, E and F, and Supplemental Figure 9, A-C). In addition, we assessed the effects of CECR2 deletion in 4T1 cells on macrophage polarization by immunofluorescence (IF) staining of the lung metastases from wild type BALB/c mice implanted with CECR2 knockout or control 4T1 cells via tail vein. Consistently, CECR2 knockout decreased M2 macrophages in metastatic tumor immune microenvironment (Supplemental Figure 9D).

**Figure 6.**
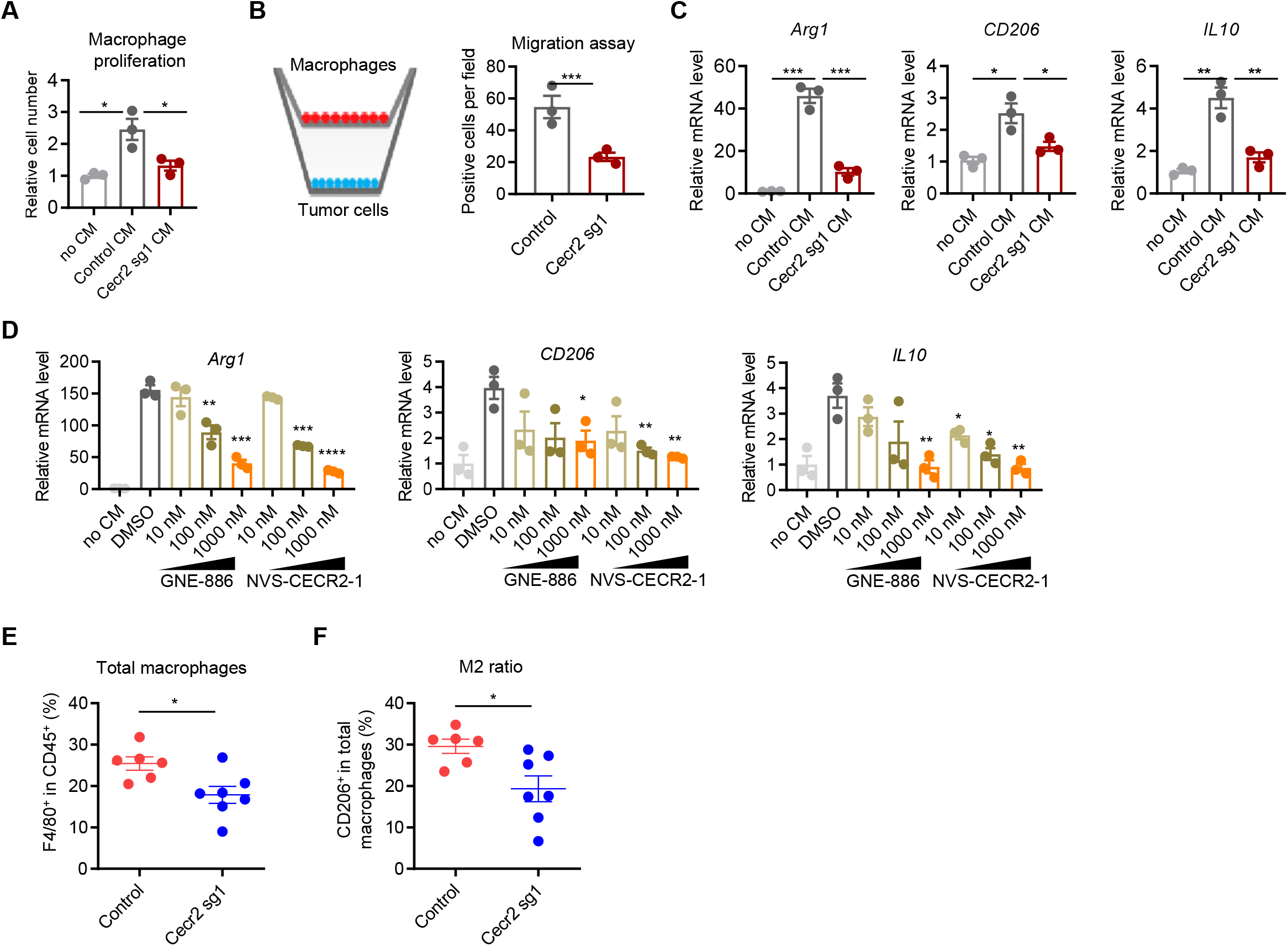
CECR2 expression in breast cancer cells increases M2 macrophages in tumor microenvironment. (**A**) CCK8 cell proliferation assays of macrophages cultured in RPMI-1640 medium with or without conditioned medium (CM) from control or Cecr2 knockout 4T1 cells. (**B**) Macrophages were seeded into the top chamber (transwell size: 8 μm), and control or Cecr2 knockout (Cecr2 sg1) 4T1 cells were seeded into the bottom chamber. Shown are schematics of transwell co-culture experiments (left panel) and quantification of migrated macrophages (right panel). (**C**) RT-qPCR analysis of M2 markers *Arg1, CD206 and IL-10* in macrophages cultured with or without condition medium (CM) from control or Cecr2 knockout 4T1 cells. (**D**) Macrophages were seeded into 6-well plate and treated with conditioned media (CM) harvested from 4T1 cells treated with DMSO, GNE-886 and NVS-CECR2-1 at indicated dosage for 2 days. RT-qPCR analyses of M2 markers *Arg1, CD206 and IL-10* in macrophages were shown. (**E-F**) Flow cytometry analyses of TAMs in the lungs from immunodeficient BALB/c nude mice with tail vein injection of control (n=6) and Cecr2 knockout (sg1) 4T1 cells (n=7) at week 2. Shown are the percentages of total macrophages (**E**) and the ratios of M2 macrophages (**F**). \**p* < 0.05; ** *p* < 0.01; *** *p* < 0.001; **** *p* < 0.0001. Representative data from triplicate experiments are shown, and error bars represent SEM.

CSF1 was shown to play major roles in regulation of macrophages (54,55). To determine if CSF1 mediates the effects of CECR2 on macrophage and tumor growth, we overexpressed CSF1 in Cecr2 knockout 4T1 tumor cells (Supplemental Figure 10A). The 4T1 cell lines with control, Cecr2 knockout (Cecr2 sg1) or Cecr2 knockout with CSF1 overexpression (Cecr2 sg1+CSF1) were injected into BALB/c mice through tail vein. The metastatic activity of those cells was assayed with India ink staining of the whole lung and H&E staining of the lung sections. These results showed that decreased lung metastasis caused by Cecr2 loss is mostly restored by CSF1 overexpression (Figure 7, A-D). We then examined the macrophage and activated CD8^+^ T cell populations in lung lesions using flow cytometry assays. We found that Cecr2 depletion in 4T1 cells strongly decreased the number of macrophages and the percentage of M2 macrophages, and increased activated CD8^+^ T cells in lung metastases, while overexpression of CSF1 suppressed these phenotypes (Figure 7, E-G, and Supplemental Figure 10, B-D). To assess the therapeutic potential of CECR2-targeted therapy *in vivo*, wild type BALB/c mice implanted with 4T1 cells via tail vein were treated with NVS-CECR2-1 or PBS every other day for 28 days (Figure 7H). We found that NVS-CECR2-1 treatment significantly inhibited the ability of 4T1 cells to metastasize to lung (Figure 7, I-K). Taken together, these results showed that CECR2 targeting inhibits macrophage polarization and breast cancer metastasis to lung.

**Figure 7.**
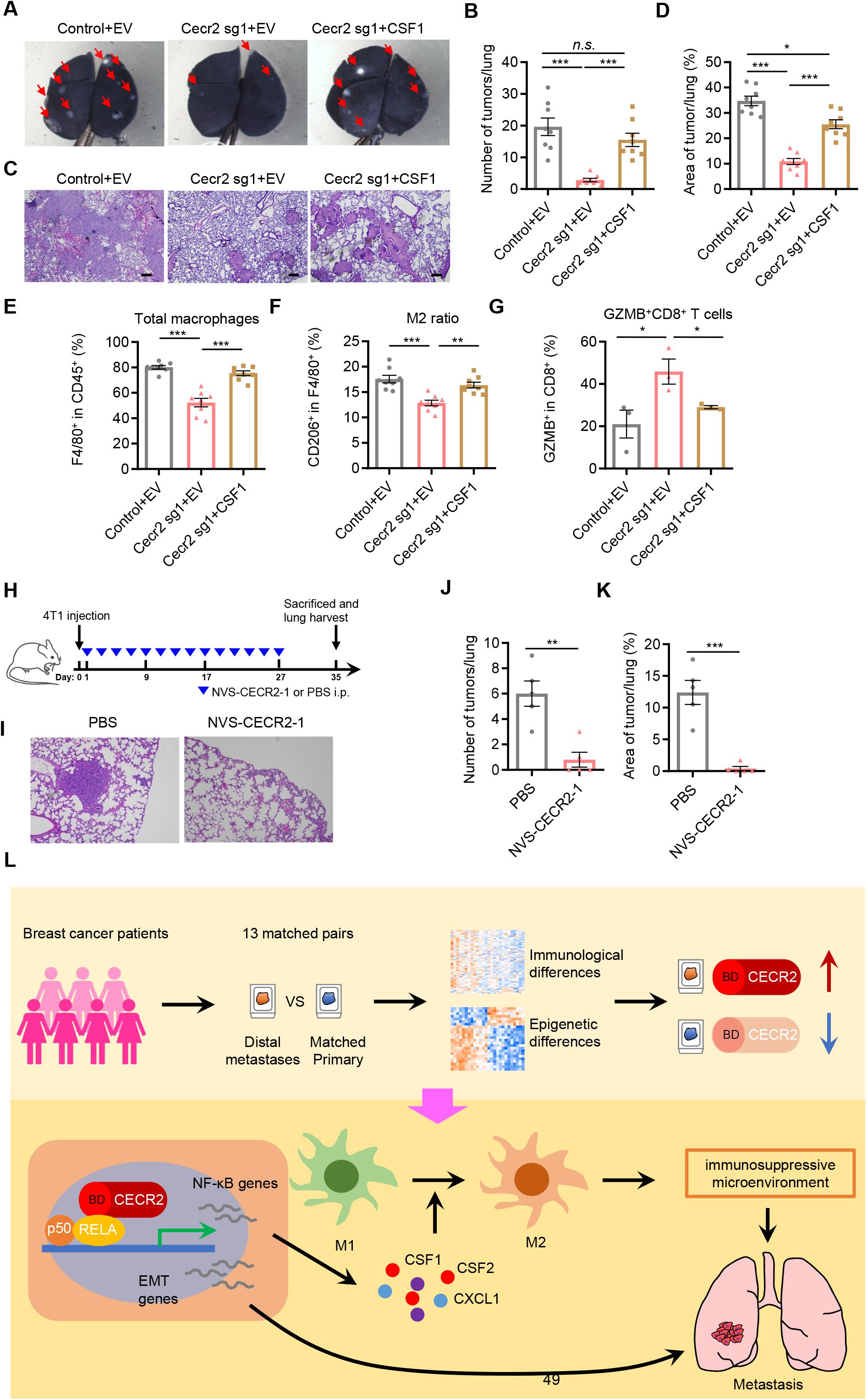
CECR2 promotes breast cancer metastasis through CSF1-mediated macrophage polarization and suppression of anti-tumor immunity. (**A-B**) BALB/c wild type mice were injected with control 4T1, Cecr2 knockout (sg1) 4T1 cells, or Cecr2 knockout 4T1 cells with CSF1 overexpression (n=8 for all the groups) through tail vein. Metastatic lesions in the lungs at week 3 after tumor cell injection were stained by India ink. Shown are representative images (**A**) and quantification of metastases in the lungs (**B**). (**C-D**) H&E staining of the lungs from mice in (A) at week 3. Shown are representative images (**C**) and quantification of tumor areas in the lungs (**D**). Scale bars: 200 μm. (**E-G**) Flow cytometry analysis of lung lesions from BALB/c wild type mice injected with control 4T1, Cecr2 knockout (sg1) 4T1 cells, or Cecr2 knockout 4T1 cells with CSF1 overexpression (n=8 for (E-F), n=3 for (G)) through tail vein at week 3. Shown are quantification of the percentages of total macrophages (CD45+F4/80+) (**E**), M2 macrophages (CD45+F4/80+CD206+) (**F**) and Granzyme B (GZMB)+ CD8+ T cells (CD45+CD8+GZMB+) (**G**). GZMB, Granzyme B. * *p*<0.05; \*\*\**p*<0.001, *n.s.* not significant. Representative data from triplicate experiments are shown, and error bars represent SEM. (**H**) Schematic illustration of intraperitoneal injection (i.p.) of NVS-CECR2-1 (10 μg/injection/mouse) or equal volume of PBS every other day for 28 days one day after tail vein injection of 4T1 cells (1×105/mouse) in BALB/c mice. All mice were sacrificed on day 35 to collect lungs and H&E staining were performed. (**I-K**) Representative H&E staining (**I**), quantification of total tumor lesions per lung (**J**) and percentage of tumor area per lung (**K**) of lungs from BALB/c mice in (**H**). (**L**) Graphical model of CECR2 promotes breast cancer metastasis by binding to RELA through its bromodomain (BD) and activating NF-κB response genes including CSF1 to modulate the immune suppressive microenvironment at the metastatic site.

## Discussion

In this study, we identified a targetable epigenetic factor CECR2 that controls metastasis by promoting M2 macrophage polarization to create an immunosuppressive microenvironment. In metastatic breast cancer cells, CECR2 interacts with acetylated RELA to activate NF-κB targets, such as CSF1, CSF2 and CXCL1. Depletion or inhibition of CECR2 suppresses NF-κB signaling and inhibits the secretion of these cytokines by tumor cells, which results in decreases of M2 macrophages. As the result, CECR2 depletion or inhibition enhances anti-tumor immunity and inhibits breast cancer metastasis (Figure 7L). These results indicate that CECR2 regulates tumor immune microenvironment to promote metastasis.

Epigenetic aberrations contribute to the initiation and maintenance of an immunosuppressive microenvironment that promotes tumor evasion (56–58). Understanding of epigenetic mechanisms controlling the immunosuppressive microenvironment, therefore, is essential for the development of epigenetic drugs to target both tumor cells and their immune microenvironments (58,59). Previous studies have shown that EZH2 and DNMT1 repress chemokines CXCL9 and CXCL10, critical for T helper 1 cell trafficking to ovarian tumors (60). Polycomb Repressive Complex 2 (PRC2)-mediated epigenetic silencing in tumor cells not only play an oncogenic role, but also contribute to blockade of CD4 and CD8 T cell recruitment into human colon cancer tissue (61). Melanoma cells overexpress H3K27 demethylase KDM6B to activate NF-κB and BMP-mediated STC1 and CCL2 expression, leading to a favorable microenvironment for melanoma growth and metastasis (62). KDM5 histone demethylases contributes to immunosuppressive microenvironment by suppression of STING in breast cancer (50), and KDM5A was shown to be critical to breast cancer metastasis (63). Here, we demonstrate the epigenetic reader CECR2 is required for metastatic breast cancer cells to express NF-κB target immune genes, including CSF1 and CXCL1, which promote an immunosuppressive microenvironment. Therefore, targeting CECR2 suppresses breast cancer metastasis partly by enhancing anti-tumor immunity.

Immune microenvironment could be conditioned actively by tumor cells to develop a permissive and supportive metastatic niche (64). We have previously found that tumor infiltrating lymphocyte (TIL) and PD-L1 protein expression is downregulated in metastatic breast tumor, as well as the key immune tolerance genes (14), which is consistent with our current analysis. Moreover, we found that tumor-associated macrophages are a major component of tumor immune microenvironment, and significantly modulate anti-tumor immunity to promote breast cancer metastasis. In breast cancer, neutrophils are recruited by factors from the primary tumor to generate the lung pre-metastatic niche, which inhibits anti-tumor CD8^+^ T cells to form an immune suppressive environment (65,66). On the other hand, patrolling monocytes are found to inhibit cancer cell metastasis by preventing cancer cell seeding in the pre-metastatic niche (67,68). The inflammatory monocytes are recruited to pre-metastatic microenvironment to facilitate breast cancer metastasis (69). Tumor-associated macrophages also promote the formation of pre-metastatic niche for cancer metastasis (70,71). Besides the contribution to pre-metastatic niche formation, several types of recruited immune cells were found to support the metastatic tumor growth. In breast cancer, neutrophils infiltrate the liver metastatic site to enhance breast cancer cell growth and metastasis (72). Macrophages polarize from a potentially tumor-inhibiting state to a tumor-promoting state in tumor microenvironment (73). In breast cancer, CSF1 was suggested to selectively promote lung metastasis by regulating the infiltration and function of tumor-associated macrophages in the PyMT breast cancer model (74). VEGFR1 signaling in metastasis-associated macrophages is crucial for breast cancer metastasis through regulating a set of inflammatory response genes, including CSF1, and CSF1-mediated autocrine signaling play a key role in tumor-promoting capability of these macrophages (75). In addition, CXCL1 produced by tumor-associated macrophages also promotes breast cancer metastasis (76,77).

Our work identifies that cytokines regulated by an epigenetic regulator CECR2, including CSF1 and CXCL1, modulate polarization and proliferation of tumor-promoting M2 TAMs. Macrophages, one of the dominant leukocytes in the tumor microenvironment of solid tumors, play essential roles in driving tumor initiation, progression and metastasis (78,79). CSF1 and CXCL1 are the major targets of CECR2, and those cytokines function in a paracrine fashion to recruit M2 TAMs to promote tumor progression and metastasis (74). M2 TAMs support cancer cells to metastasize to distant organs (80). Moreover, they express molecular triggers of checkpoint proteins that suppress T-cell activation (81). Consistently, we find that CECR2 depletion reversed immune suppression at lung metastatic sites in breast cancer, suggesting that CECR2 promotes an immunosuppressive microenvironment at the metastatic sites. These results also suggest clinically testable therapeutic strategies.

Bromodomain is the acetyl lysine ‘reader’ module in epigenetic factors, and targeting bromodomain has been shown to promote anti-inflammatory and anti-cancer activities (82). Multiple inhibitors against bromodomain and extra-terminal domain (BET) proteins are already in clinical testing (79). Similar to BET bromodomains, the bromodomain of CECR2 is predicted to be highly druggable (28). Indeed, pharmacological inhibitors of CECR2 NVS-CECR2-1 and GNE-886 have been developed. In fact, treatment with these CECR2 inhibitors substantially suppressed the expression of CECR2 targets CSF1 and CXCL1 in multiple metastatic cancer cells, suggesting a possible therapeutic approach to inhibit immunosuppression in the metastatic tumor microenvironment. Our results also support testing of anti-CSF1 therapeutic antibodies (MCS110, PD-0360324) in the clinic. We consider CECR2 bromodomain inhibition as a promising novel therapeutic strategy to treat metastatic breast cancer. This strategy reduces immune suppression at the metastatic sites and might increase the efficacy of immunotherapies.

## Methods

### Plasmids, compounds, and cell culture

GFP-CECR2 (Addgene, #65385) was transferred into pDONR221 by Gateway BP Clonase II enzyme mix (Thermo Fisher, # 11789020,), and bromodomain was deleted (ΔBRD) by mutagenesis. Both CECR2 wildtype (WT) and ΔBRD donor plasmids were then transferred to pMH-SFB (Addgene, #99391) by Gateway LR Clonase II Enzyme Mix (#11791020, Thermo Fisher) to generate pMH-SFB-CECR2 (FLAG-CECR2). RelA cFlag pcDNA3 (FLAG-RELA, addgene, #20012), T7-RELA (Addgene, # 21984), T7-RELA-K310R (Addgene, #23250) were obtained from Addgene. NVS-CECR2-1 (SML-1803) was purchased from Sigma, St Louis, MO, and GNE-886 was obtained from Genentech, South San Francisco, CA. 4T1 breast cancer cells were cultured in RPMI1640 supplemented with 10% fetal bovine serum, 100 U/mL penicillin, and 100 μg/mL streptomycin. MDA-MB-231(MDA231), MDA231 LM2 (LM2), MDA231 BoM (BoM) and MDA231 BrM2 (BrM2) breast cancer cells and HEK293T cells were cultured in Dulbecco’s Modified Eagle Medium supplemented with 10% fetal bovine serum and 100 U/mL penicillin, and 100 μg/mL streptomycin. Cells were periodically tested for mycoplasma contamination and authenticated using short tandem repeat profiling.

Knockout sgRNAs were designed according to online software CHOPCHOP (https://chopchop.rc.fas.harvard.edu/) and cloned into LentiCRISPRv2 vector. CECR2/Cecr2 knockout LM2 and 4T1 cells were generated as described previously (83). Briefly, 1.5 μg lentiviral plasmid, 1 μg psPAX2, and 0.5 μg pMD2.G were transfected into HEK293T cells in 6-well plates by Lipofectamine 2000 Transfection Reagent (Invitrogen) according to the manufacturer’s instructions. Fresh growth medium was replaced on the following day. Then after 48 hours, lentivirus-containing media were harvested and filtered through a 0.45 μm filter. Target cells were infected with lentiviruses, and fresh growth medium was then refed to cells after 24 hours. After 48 hours of medium change, cells were selected with 2 μg/ml puromycin for 1-2 weeks for stable knockout cell lines. sgRNA controls were described previously (84). Primers for knockout were human CECR2 sg1: TGATGTCCTCTAGGTAGCTG; human CECR2 sg2: CGCTCTTCACAGAGATGACG; mouse Cecr2 sg1: GAGTACGCAGAGGAAGGTCT; mouse Cecr2 sg2: GAGATGTGCCCGGAGGAAGG; human RELA sg: AGACGATCGTCACCGGATTG. For knockout detection, one primer is using the sg sequence and the other primer were gcecr2-sg1: GAGTACGCAGAGGAAGGTCT; gcecr2-sg2: TCGATCTCGAAGTCGGGC. For human CECR2 or CSF1 reconstitution expression, 4T1 cells with Cecr2 knockout were transfected with human CECR2 expression vector (GFP-CECR2) or CSF1 plasmid (Obio Technology Shanghai Corp., Ltd, China, #m13002), respectively, with X-tremeGENE HP DNA Transfection Reagent (Roche, #06366236001) according to the manufacturer’s instructions, and selected with 10 μg/ml blasticidin for 2 weeks.

### Western blot and Co-immunoprecipitation (Co-IP)

Cells were washed with PBS and lysed in high salt buffer (50 mM Tris-HCl pH 7.6, 320 mM NaCl, 0.1 mM EDTA, 0.5% NP-40) or RIPA buffer (50 mM Tris-HCl pH 7.4, 150 mM NaCl, 1 mM EDTA, 1% Triton X-100, 1% sodium deoxycholate, 0.1% SDS) supplemented with protease inhibitors (PI, Roche). For nuclear and cytoplasmic extraction, NE-PER™ Nuclear and Cytoplasmic Extraction Reagents kit (#78833, Thermo Fisher) was used according to the manufacturer’s instructions. Protein concentrations were then determined by Bradford Assay (Bio-Rad Laboratories, Inc.). Samples were then boiled, resolved in SDS-PAGE, and blotted with the primary and secondary antibodies as described (84).

For exogenous Co-IP experiments, HEK293T cells were transfected with T7-RELA, T7-RELA-K310R, GFP-CECR2, FLAG-CECR2(WT and ΔBRD), FLAG-RELA plasmids as indicated with X-tremeGENE HP DNA Transfection Reagent. After 48 hours, the cells were lysed with high salt buffer (50 mM Tris-HCl pH 7.6, 320 mM NaCl, 0.1 mM EDTA, 0.5% NP-40) including protease inhibitor cocktail (Roche) on ice. For endogenous Co-IP experiments, LM2 and 4T1 cells were collected for protein extraction with high salt buffer. The prepared protein extracts were incubated with antibodies as indicated for overnight at 4 °C, followed by incubation with protein A/G beads (Pierce, #20421) for 2 hours at 4 °C for the immunoprecipitation and western blot assays.

The following antibodies were obtained commercially: rabbit anti-CECR2 (HPA002943), mouse anti-FLAG (M2, F1804), and mouse anti-tubulin (T5168) (Sigma, St. Louis, MO); mouse anti-CECR2 (C3, sc-514878), mouse anti-CSF1 (D4, sc-365779), mouse anti-NF-κB p50 (E-10, sc-8414) (Santa Cruz, Dallas, TX); rabbit anti-NF-κB p65 (D14E12, #8242), rabbit anti-NF-κB2 p100/p52 (#4882), rabbit anti-RelB (C1E4, #4922), rabbit anti-c-Rel (D4Y6M, #12707), mouse anti-GAPDH (D4C6R, #97166) (Cell Signaling Technology, Danvers, MA); rabbit anti-H3(ab1791), mouse anti-RNA pol II (8WG16, ab817) (Abcam, Cambridge, UK); rabbit anti-T7 (AB3790) and rabbit anti-H3K9/18Ac (07-593) (Millipore sigma, Burlington, MA).

### RT-qPCR and ChIP-qPCR analyses

For RT-qPCR assays, total RNA was extracted by RNeasy Mini Plus kit (Qiagen) and reverse transcription was performed using SuperScript™ III First-Strand Synthesis System (#18080051, Thermo Fisher Scientific). For one real-time PCR reaction, cDNA corresponding to approximately 10 ng of starting RNA was used and qPCR was performed with SYBR green master mix (Bio-Rad Laboratories, Inc.). Primers for real-time PCR were GAPDH-F: TGCACCACCAACTGCTTAGC, GAPDH-R: GGCATGGACTGTGGTCATGAG; CECR2-F: GCATTTGCCATCTTCTCCAT, CECR2-R: TTCCCATTCTCCACGATCTC; CSF1-F: TGGCGA GCAGGAGTATCAC, CSF1-R: AGGTCTCCATCTGACTGTCAAT; CSF2-F: TCCTGAACCTGAGTAGAGACAC, CSF2-R: TGCTGCTTGTAGTGGCTGG; CXCL1-F: ATTCACCCCAAGAACATCCA, CXCL1-R: CACCAGTGAGCTTCCTCCTC; Hprt1-F: CATAACCTGGTTCATCATCGC, Hprt1-R: TCCTCCTCAGACCGCTTTT; Gapdh-F: TTGATGGCAACAATCTCCAC, Gapdh-R: CGTCCCGTAGACAAAATGGT; Csf1-F: GTGTCAGAACACTGTAGCCAC, Csf1-R: TCAAAGGCAATCTGGCATGAAG; Csf2-F: GGCCTTGGAAGCATGTAGAGG, Csf2-R: GGAGAACTCGTTAGAGACGACTT; Cxcl1-F: ACTGCACCCAAACCGAAGTC, Cxcl1-R: TGGGGACACCTTTTAGCATCTT.

The ChIP-qPCR assays were conducted as described previously (85). Briefly, 1×10^7^ LM2 (control and knockout) and 4T1 (control and knockout) cells were cultured for each IP. Crosslinking was performed with 1% formaldehyde in culture media for 10 min, and then stopped by addition of 0.125 M glycine for 10 min. After washing, cells were collected and resuspended in the lysis buffer 1 (50 mM Hepes pH 7.5, 140 mM NaCl, 1 mM EDTA, 10% glycerol, 0.5% NP40, 0.25% Triton X-100) with complete protease inhibitor cocktail (Roche Molecular Biochemicals) for incubation on ice for 20 min. Then the cellular nuclei were spin down and resuspended in lysis buffer 2 (10 mM Tris·HCl pH 8.0, 200 mM NaCl, 1 mM EDTA, 0.5 mM EGTA) with complete protease inhibitor cocktail. After rocking for 10 min, nuclei were spin down and resuspended in lysis buffer 3 (10 mM Tris pH 8.0, 1 mM EDTA, 0.5 mM EGTA) with protease inhibitor cocktail. Then, sonication was performed to fragment chromatin to an average length of 0.5 kb. After the pre-clearance with 50 μl protein A/G agarose beads for each IP, the target or control IgG antibody was added and incubated at 4°C overnight. Then, 60 μl protein A or G agarose beads were added and incubated at 4°C for 2 hours for immunoprecipitation. The immunocomplexes were then eluted from the agarose beads and incubated at 65°C overnight to reverse crosslinking. DNA from ChIP and input were then purified for qPCR with SYBR green master mix (Bio-Rad Laboratories, Inc.). Primers for ChIP-qPCR were CSF1-F: TTGGGACGATCATAGAGCGC; CSF1-R: GTCACCCTCTGTCTTCTGCG; CXCL1-F: CTGCTGCTCCTGCTCCTG; CXCL1-R: CTGACTGAGCGAGGCTGTC; Csf1-F: GGGGCATGTGGTTTATGGGA; Csf1-R: ACTTTGAGGAGGCTGCACAG; Cxcl1-1 F: ACAGCTTTCCCGTGGACTTT; Cxcl1-R: CAGGGAGGCATGTGAAGAGG.

### Colony formation, WST1, migration and invasion assays

Colony formation assays were done by seeding single cells in 6 well plates. Colonies were fixed with 4% paraformaldehyde (PFA) (#28908, Thermo Fisher), followed by crystal violet staining for 0.5 hour. For WST1 cell proliferation assays (#11644807001, Roche), cells were seeded in 96 well plate for indicated days growth, and then were assayed according to the manufacturer’s instructions. For migration and invasion assays, tumor cells were starved in medium containing 0.2% FBS for overnight. Then, tumor cells were seeded into trans-well inserts or matrigel coated trans-well inserts with 8 μm pores (BD Biosciences), using 10% FBS as a chemoattractant. After 6 or 18 hours, trans-wells were cleaned and fixed in 4% paraformaldehyde. Cells on the apical side of each insert were scraped off and the cells on the trans-well membrane were counterstained with DAPI. Migrated and invaded cells were visualized with Keyence BZ-X700 immunofluorescent microscope. Three random fields of pictures of each three replicates were captured for quantification using ImageJ software (NIH).

### Animal studies

Female Athymic Nude-*Foxn1^nu^* immunodeficient (6-8 weeks old) mice (Envigo) were used for lung-metastasis experiments with human cell lines. The viable CECR2 knockout and control LM2 cells (3×10^5^) were re-suspended in 0.1 ml saline and injected into mice through the tail vein. For 4T1 cells, the indicated Cecr2 knockout, Cecr2 knockout with CECR2 reconstitution expression or control cells (2×10^5^) were resuspended in 0.1 ml saline, and then injected into the tail vein of female BALB/c mouse (6-8 weeks old). The 4T1 Cecr2 knockout and control cells (1×10^5^) were resuspended in 0.1 ml saline, and then injected into the tail vein of female BALB/c nude mouse (6-8 weeks old). The bioluminescence signal of lung-metastatic colonization was monitored with a Xenogen IVIS system coupled to Living Image acquisition and analysis software (Xenogen), and the signal were then quantified at the indicated time points as previously described (48). Values of luminescence photon flux of each time point were normalized to the value obtained immediately after xenografting (day 0).

For mammary fat pad tumor assays, the Cecr2 knockout and control 4T1 cells (5×10^4^) were resuspended in 0.1 ml saline, and then injected into mammary fat pad (the 4^th^ mammary glands) of BALB/c mouse (6-8 weeks old). Tumor were monitored every 3 days by measuring the tumor length (L) and width (W). Tumor volume was calculated as V=L×W^2^/2. Mice were euthanized when primary tumors reached 1,000 mm^3^. The lungs were harvested for hematoxylin and eosin (H&E) staining. In the CECR2 inhibitor treatment experiment, 4T1 cells (1×10^5^) were injected into each mice through tail vein. NVS-CECR2-1 (10 μg/injection/mouse) or equal volume of PBS was injected into mice by intraperitoneal injection every other day for 28 days. All mice were sacrificed on day 35 to collect lungs and H&E staining were performed. All animal procedures were approved by the Institutional Animal Care and Use Committee of Yale University and Central South University.

### Histopathology

Mice were euthanized by CO_2_ asphyxiation and lungs were harvested, immersion-fixed in 10% neutral buffered formalin, processed, sectioned and stained by hematoxylin and eosin (H&E) with routine methods by Yale Research Histology (Department of Pathology) or Comparative Pathology Research (Department of Comparative Medicine). Tissues were evaluated blindly to experimental manipulation for the presence and number of tumor metastatic foci and percentage of lung effaced by tumor. Digital light microscopic images were recorded using an Axio Imager.A.1 microscope and an AxioCam MRc5 camera and AxioVision 4.7 imaging software (Zeiss) and optimized in Adobe Photoshop (Adobe Systems Incorporated, USA), or using a Keyence BZ-X700 immunofluorescent microscope.

### Immunofluorescence (IF) and immunohistochemistry (IHC) staining

For IF staining of cells, cells were seeded on coverslips, fixed with 4% paraformaldehyde for 10 minutes, permeabilized with 0.4% Triton X-100 in PBS for 5 minutes, and then blocked with 10% NGS (Normal goat serum) before incubation with primary antibodies at 4°C for overnight. For IF and IHC staining of paraffin embedded tissue, all samples were sectioned and deparaffinized with xylene, followed with ethanol washing, antigen retrieval by heat in EDTA buffer (PH 8.0) or citrate buffer (PH 6.0), tissue samples were penetrated by methanol and blocked with BSA before incubation with primary antibodies at 4°C overnight. For IF staining, second antibodies were applied, which was followed with DAPI staining. For IHC staining, DAB reaction and hematoxylin staining were used. All the stained samples were visualized with a Keyence BZ-X700 immunofluorescent microscope at 4X, 10X and 20X. Three random fields of pictures of each three replicates were captured for quantification using ImageJ software (NIH).

### Preparation of coeliac macrophage, conditioned medium and co-culture

The 3% thioglycollate broth was injected into mouse abdomen and macrophages were harvested and purified 3 days later. To get tumor conditioned medium (CM), tumor cells were grown to 50% and then changed to 2% FBS culture medium for 3 days. CM was then collected, concentrated and frozen at −80°C for long term use. For co-culture assays, macrophages were seeded at upper chamber and tumor cells were seeded at lower chamber. After 12 hours, migrated macrophages were stained and counted.

### Flow cytometry analyses

Cells were prepared for single cell suspension and were fixed with 2% paraformaldehyde solution in PBS. After being washed with a flow cytometry staining buffer, cells were stained with fluorescent-labeled antibodies for cell-surface markers for 1 hour on ice in the dark. The cells were then washed and resuspended in the flow cytometry staining buffer for flow cytometry analysis. The follow antibodies were used: anti-mouse F4/80 PE (123109), anti-mouse F4/80 APC (100311), anti-mouse CD11b FITC (101205), anti-mouse CD206 PE (141705), anti-mouse CD45 APC (103112), anti-mouse CD8 PE (123110), anti-mouse F4/80 PE (123110), anti-mouse CD45 APC-Cy7 (110716), anti-mouse CD86 PerCP-Cy5.5 (105028), anti-mouse CD206 APC (141708), anti-mouse/human Granzyme B PE (372207), anti-mouse/human Granzyme B AF647 (515405) (BioLegend, San Diego, CA). Flow cytometry was performed on a LSRII flow cytometer or FACSCalibur (BD Biosciences). Data were analyzed with FlowJo or BD CellQuest Pro software version 5.1.

### India INK staining

The animal was placed on its back after being euthanized by CO_2_. The rib cage was cut open to expose the lungs and an incision was made on the neck to expose its trachea carefully. 2 ml of India ink solution (85% India ink / 15% ddH_2_O) was slowly infused into the lungs through the trachea by a 25-gauge needle. The infused lung samples were kept in Fekete’s solution (900 mL 70% ethanol / 90 mL 37% formaldehyde / 15 mL 91% acetic acid) for de-staining. The tumor nodules do not absorb India ink, which results in the normal lung tissue staining black while the tumor nodules remain white. White tumor nodules were counted blindly by 3 individuals and the numbers were recorded and averaged as the tumor count on the lungs for each of the animals. Lung samples were then further processed for the H&E staining to look for micro-metastases inside the lungs.

### RNA-sequencing and bioinformatics analysis

FFPE RNA was extracted from matched primary and metastatic FFPE samples by QIAGEN AllPrep DNA/RNA FFPE kit. RNA of control and knockout LM2 cells was isolated using RNeasy Plus Mini Kit (Qiagen, Hilden, Germany). All the patient FFPE sample libraries are prepared with TruSeq RNA Access Library Prep Kit (Illumina, #RS-301-2001). All the cell line mRNA libraries for sequencing were prepared according to the TruSeq Stranded Total RNA Library Prep Kit (Illumina, #RS-122-2201). Sequencing (75 bp, paired end) was performed using Illumina HiSeq 2000 sequencing system at the Genomics Core of Yale Stem Cell Center or Illumina HiSeq 2500 sequencing system at Yale Center for Genome Analysis (YCGA). RNA-seq data were deposited in the National Center for Biotechnology Information (NCBI) Gene Expression Omnibus database under GSE148005 (https://www.ncbi.nlm.nih.gov/geo/query/acc.cgi?acc=GSE148005, with secure token uhedgyskrxgdnwh).

The RNA-seq reads were mapped to human genome (hg19 for tumors or hg38 for cell lines) with Bowtie2 (86,87). The uniquely mapped reads (only keep alignment with MAPQ >=10) were counted to ENCODE gene annotation (version 24) using FeatureCounts (87,88). Differential gene expression was performed with DESeq2 (89). Gene expression values were then transformed by variance-stabilizing transformation (VST) with DESeq2 and batch effects were removed using ComBat (90). After normalizing each gene to Z-score, heatmap were then plotted with heatmap2 (91). Gene expression profiles of control or knockout cells were used for Gene Set Enrichment Analysis (GSEA) using GSEA version 2.0 software (92). The gene set database of h.all.v6.1.symbols.gmt (Hallmarks) was used. Statistical significance was assessed by comparing the enrichment score to enrichment results generated from 10,000 random permutations of the gene set. Kaplan-Meier Plotter analyses (https://kmplot.com/analysis/) (46) were performed for the distant metastasis free survival of breast cancer patients, relapse free survival and post progression survival of gastric and ovarian cancer patients based on the Jetset best probe set (239752_at) for CECR2 mRNA level. The following settings were used for the analysis: distant metastasis-free survival, autoselect best cutoff, 150 months follow-up threshold. Percentage of immune cells in the 13 pairs of patient samples was calculated using CYBERSORTx (https://cibersortx.stanford.edu/) (42). Deregulated epigenetic genes comparing matched metastases vs primary samples and deregulated hallmark gene sets of CECR2 knockout samples were analyzed with online tool Venny 2.1 (https://bioinfogp.cnb.csic.es/tools/venny/) to generate the Venn diagrams.

### Statistical analysis

Comparisons between two groups were performed using an unpaired two-side Student’s *t* test. Comparisons between matched data of metastasis and primary tumor samples from the sample breast cancer patient were performed using a paired Student’s *t* test. Comparisons of multiple conditions was done with one-way ANOVA and Tukey’s post hoc test. *p*<0.05 was considered significant. Graphs represent either group mean values ± SEM or individual values (as indicated in the figure legends). For animal experiments, each tumor graft was an independent sample. For correlation analysis, the Pearson coefficient was used. All experiments were reproduced at least three times.

## Supporting information

Supplemental Table of content and Figures 1-10

## Acknowledgments

We would like to thank all members of Yan, Stern and Nguyen laboratories at Yale University for helpful discussions, Lori Charette and Dr. Yalai Bai at Yale Pathology Tissue Services for TMA building, Dr. Mei Zhong at Yale Stem Cell Center Genomics Core facility for helping with sample preparation for RNA-seq, Dr. Joan Massagué at Memorial Sloan Kettering Cancer Center for providing MDA-MB-231, LM2, and 4T1 cells, Dr. Yibin Kang at Princeton University for providing BrM2 cells, Dr. Don Nguyen for providing PC9-BrM4 cells, Dr. Marcus Bosenberg for providing YUMM1.7 cells, Dr. Andrea Cochran at Genentech for providing GNE-886, and Dr. Narendra Wajapeyee at the University of Alabama Birmingham for helping with compiling the epigenetic gene list. Sequencing done at Yale Stem Cell Center Genomics Core facility was supported by the Connecticut Regenerative Medicine Research Fund and the Li Ka Shing Foundation.

## Author contributions

M.Z., M.Y. and Q.Y. designed the research. Q.Y. conceived and oversaw the project. M.Z. performed most of the experiments. M.Y., Y.Z. and M.Z. performed in vivo and in vitro assays related to macrophages. Z.Z.L. and M.Z. performed the bioinformatic analysis. M.Z., K.A., A.A., Z.T. and M.Y. performed animal studies. M.Z. performed the in vivo imaging experiments. C.J.B. performed histological analyses in Figure 2 F and G and M.Z., and M.Y. performed histological analyses in the other figures. L.H.C. cloned FLAG-CECR2 constructs. S.M.L. and Y.A. performed some cell culture work. H.S. performed some flow cytometry analysis. S.J.R., V.B., J.S.M., L.P., and D.L.R. provided clinical samples, collected clinical information, and helped with experimental design related to clinical samples. M.Z., Z.Z.L., K.A., C.J.B., X.C., M.Y., and Q.Y. analyzed the data. M.Z., M.Y. and Q.Y. wrote the paper.

