## Supplemental Table of content and Figures 1-10 for "CECR2 Drives Breast Cancer Metastasis by Suppressing Macrophage Inflammatory Responses"

**Supplemental Figure 1**. The ratios of M1 macrophages to total Macrophages is decreased in matched metastases.

**Supplemental Figure 2**. CECR2 expression negatively correlates with metastasis-related survival.

**Supplemental Figure 3**. Knockout of CECR2 inhibits invasion of human breast cancer cells without affecting their proliferation.

**Supplemental Figure 4**. Knockout of Cecr2 inhibits invasion of mouse breast cancer cells without affecting their proliferation in vitro and in vivo.

**Supplemental Figure 5**. Pathways affected by CECR2 knockout in LM2 cells.

**Supplemental Figure 6**. CECR2 interacts with RELA to activate NF-κB response genes.

**Supplemental Figure 7**. CECR2 inhibition suppresses the expression of NF-κB response genes as well as the migration and invasion capability.

**Supplemental Figure 8**. CECR2 inhibitors do not affect macrophage polarization in vitro.

**Supplemental Figure 9**. Cecr2 loss in 4T1 cells does not affect the percentages of M1-like macrophages and NK cells in lung metastases.

**Supplemental Figure 10**. CECR2 activates CSF1 to inhibit anti-tumor immunity in metastatic microenvironment.

**Supplemental Table 1.** Patient information for 13 paired breast tumor samples used for RNA-seq analysis.

**Supplemental Table 2.** Significantly differentially expressed genes in 13 pairs of matched distant metastases compared with primary tumors.

**Supplemental Table 3**. Expression of 29 immuno-oncology targets currently in clinical development in paired primary and metastatic breast cancer samples.

**Supplemental Table 4**. Percentage of immune cells by CIBERSORTx analysis of RNA seqencing data of 13 pairs of matched distant metastases and primary tumors.

**Supplemental Table 5**. Epigenetic gene (epigene) list

**Supplemental Table 6**. Significantly differentially expressed epigenetic genes in 13 pairs of matched distant metastases compared with primary tumors.

**Supplemental Table 7**. Correlation of epigenetic genes with M2 ratio in total macrophages and the hazard ratio of epigenetic genes of distant metastasis free survival.

**Supplemental Table 8**. CECR2 IHC scores of tissue microarray.

**Supplemental Table 9**. RNA sequencing analysis of CECR2 knockout-1 (sg1-1 and sg1-2) versus control (Control-1 and Control-2) LM2 cells.

**Supplemental Table 10**. RNA sequencing analysis of CECR2 knockout-2 (sg2-1 and sg2-2) versus control (Control-1 and Control-2) LM2 cells.

**Supplemental Table 11**. Negatively enriched hallmark pathways by gene set enrichment analysis (GSEA) comparing CECR2 sg1 with Control LM2 cells.

**Supplemental Table 12**. Negatively enriched hallmark pathways by gene set enrichment analysis (GSEA) comparing CECR2 sg2 with Control LM2 cells.

**Supplemental Table 13**. Positively enriched hallmark pathways by gene set enrichment analysis (GSEA) comparing CECR2 sg1 with Control LM2 cells.

**Supplemental Table 14**. Positively enriched hallmark pathways by gene set enrichment analysis (GSEA) comparing CECR2 sg2 with Control LM2 cells.

### Supplemental Figure 1

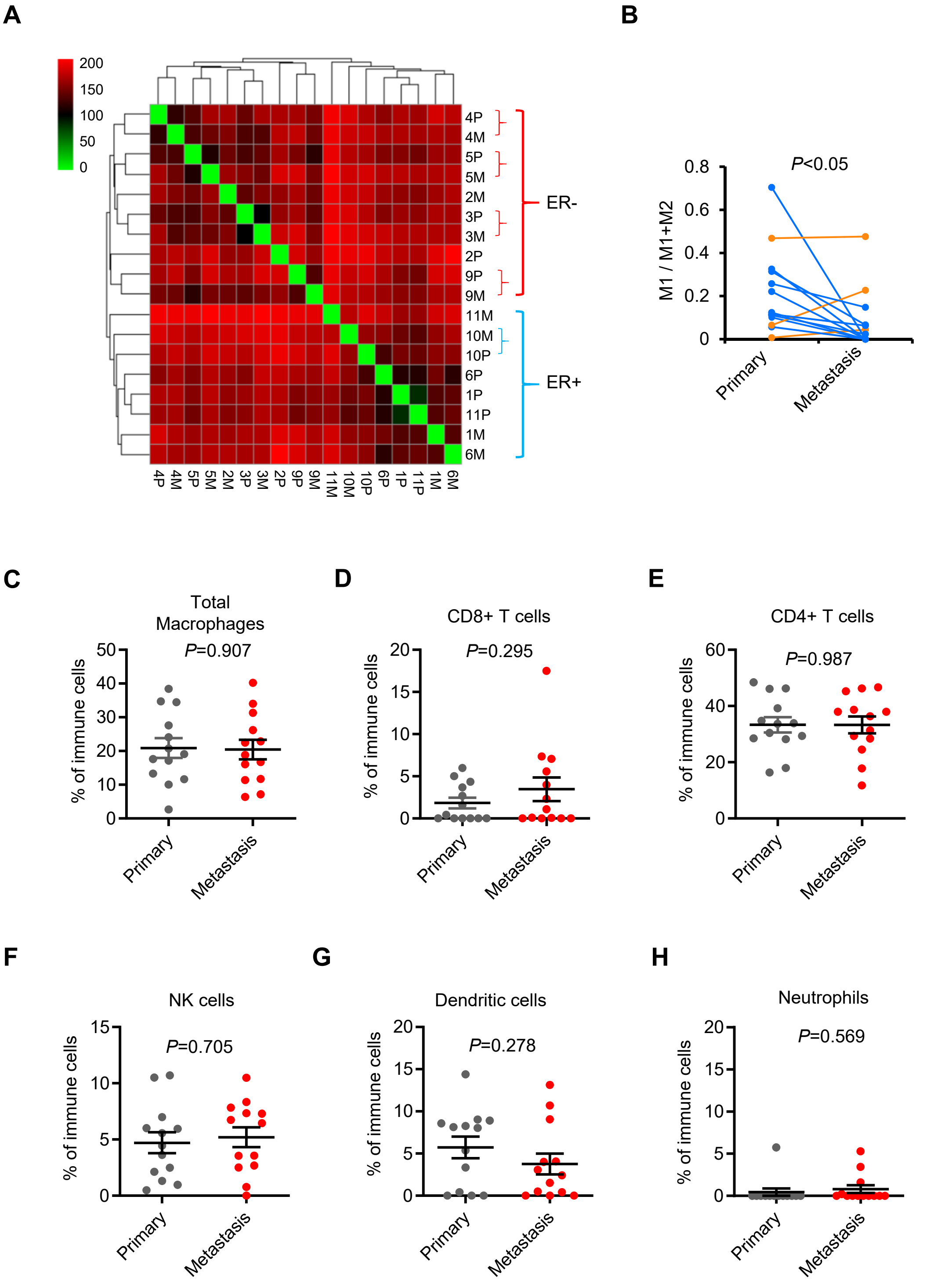


**Supplemental Figure 1. The ratios of M1 macrophages to total Macrophages is decreased in matched metastases.** (**A**) Clustering of primary breast tumors and matched metastatic samples based on their transcriptomes. (**B-H**) RNA-seq data of matched primary tumors and distal metastases from 13 breast cancer patients were analyzed by CIBERSORTx and immune cell composition of complex tissues were characterized from their gene expression profiles. The ratios of M1-like macrophages to total TAMs (**B**), percentage of total macrophages (**C**), CD8+ T cells (**D**), CD4+ T cells (**E**), NK cells (**F**), dendritic cells (**G**) and neutrophils (**H**) in immune cells. Yellow lines marks the samples with increased ratio in metastasis while blue lines marks the ones with decreased ratio.

### Supplemental Figure 2

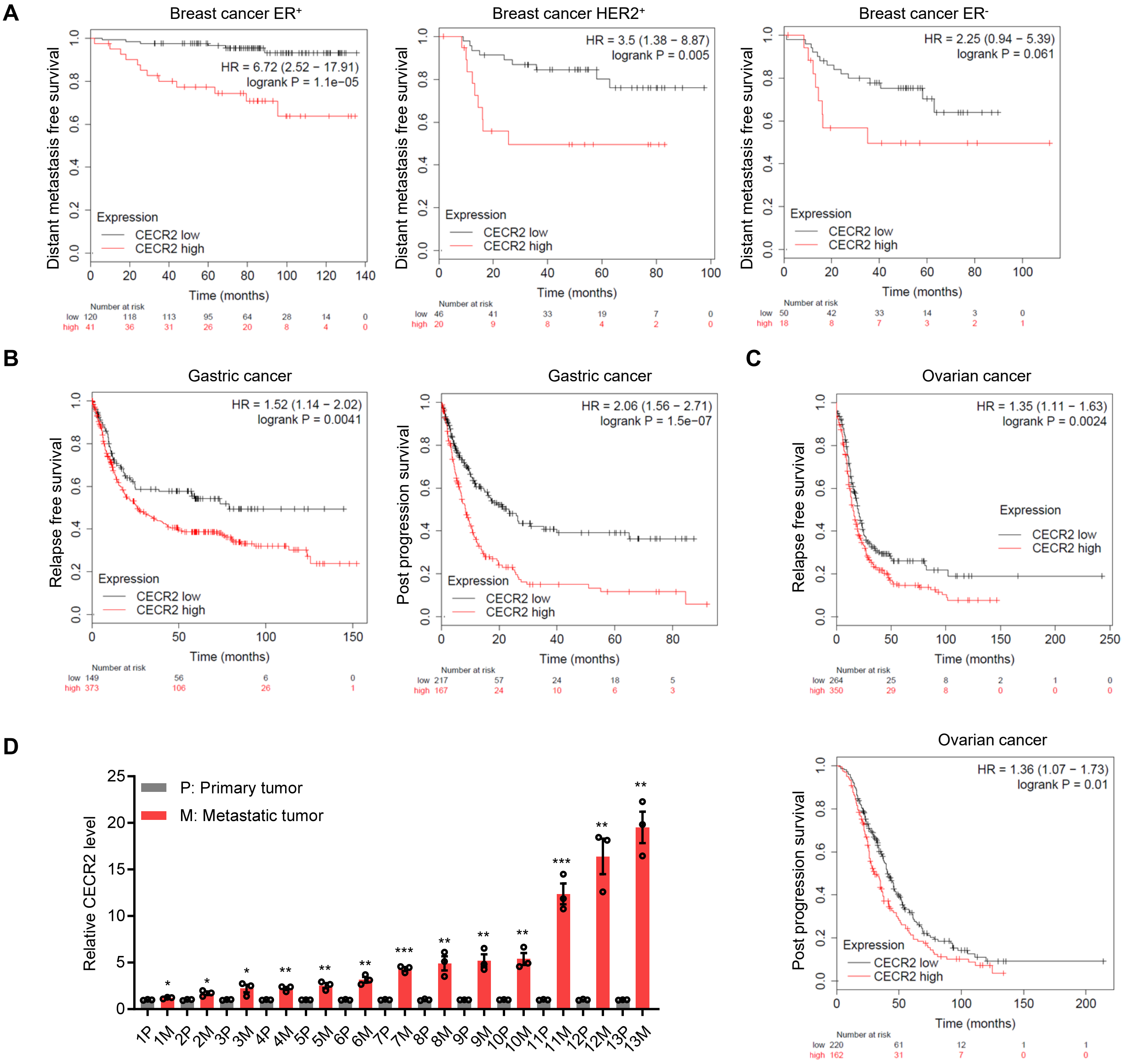


**Supplemental Figure 2. CECR2 expression negatively correlates with metastasis-related survival.**
(**A-C**) Kaplan-Meier plotter analyses showing association of CECR2 mRNA levels with distant metastasis free survival in $ER^-$, $HER2^+$ and $ER^+$ breast cancer patients (A), with relapse free survival and post progression survival in gastric cancer (**B**) and ovarian cancer (**C**). Hazard ratio (HR) and log-rank *p* values were calculated. (**D**) RT-qPCR analysis of CECR2 mRNA levels in the matched metastatic and primary tumor samples. **p* < 0.05, ** *p* < 0.01, *** *p* < 0.001. Representative data from triplicate experiments are shown, and error bars represent SEM.

#### Supplemental Figure 3

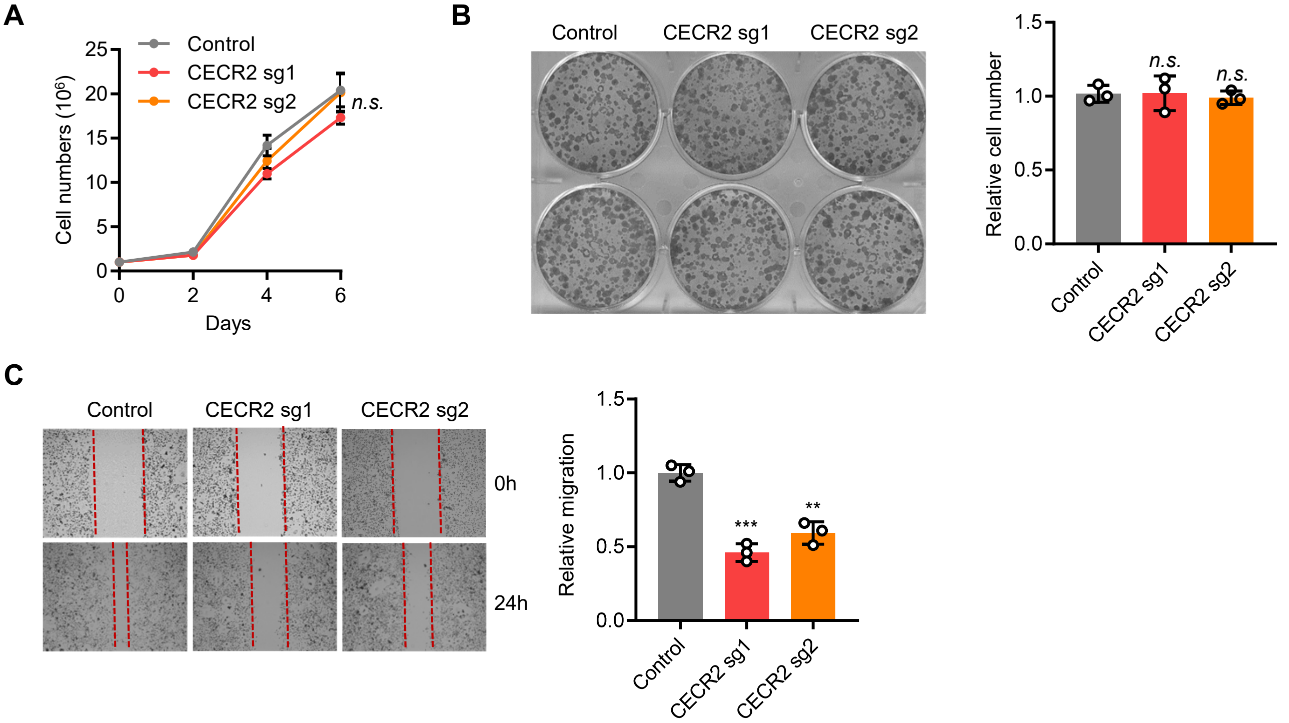


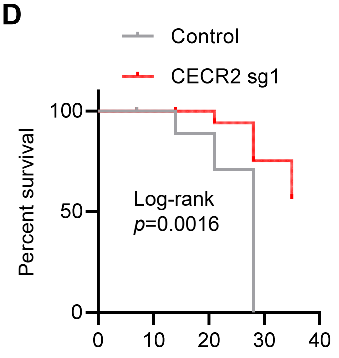


**E**

| Score | Level | % |
|---|---|---|
| 0 | none | 0 |
| 1 | minimal | <10% |
| 2 | mild | 10-25% |
| 3 | moderate | 25-50% |
| 4 | marked | 50-75% |
| 5 | severe | 75-100% |

**Supplemental Figure 3. Knockout of CECR2 inhibits invasion of human breast cancer cells without affecting their proliferation.**

(**A**) WST1 cell proliferation assays comparing control and CECR2 knockout LM2 cells. (**B**) Representative images (left panel) and quantification (right panel) of colony formation assays comparing control and CECR2 knockout LM2 cells. (**C**) Representative images at 24 hours (left panel) and quantification of the closure of wound distance (right panel) in scratch assays comparing control and CECR2 knockout LM2 cells. (**D**) Kaplan-Meier survival curve of nude mice with tail vein injection of control (n=8) or CECR2 knockout (CECR2 sg1) LM2 cells (n=7) using bioluminescence signal of $5 \times 10^6$ as the end point. Mantel-Cox log-rank test was performed to calculate the p value. (**E**) Parameters for tumor score calculation. ** $p < 0.01$, *** $p < 0.001$, *n.s.*, not significant. Representative data from triplicate experiments are shown, and error bars represent SEM.

### Supplemental Figure 4

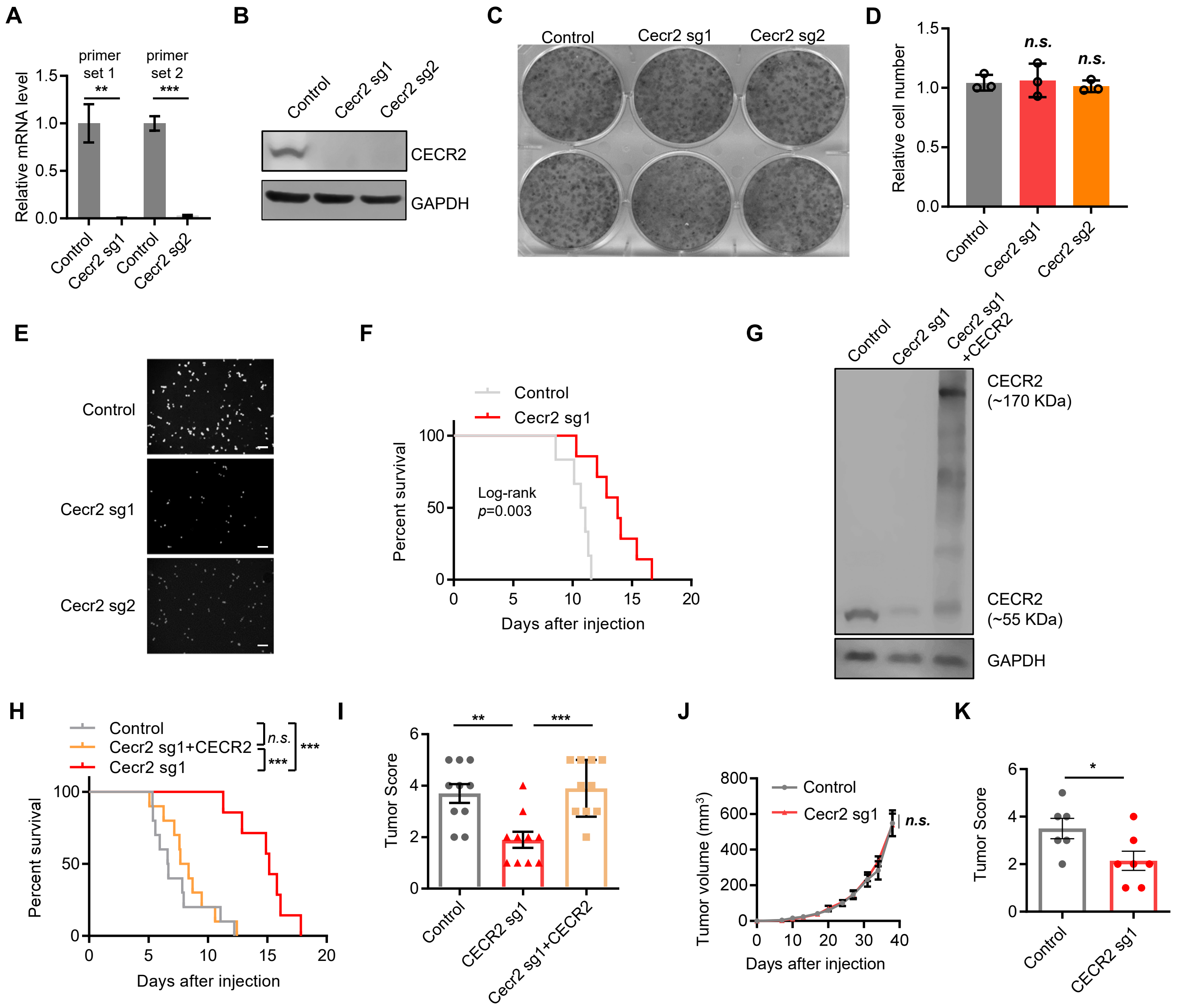


**Supplemental Figure 4. Knockout of Cecr2 inhibits invasion of mouse breast cancer cells without affecting their proliferation *in vitro* and *in vivo*.**

(**A**) Cecr2 knockout in 4T1 cells (sg1 and sg2 ) was validated using primer sets spanning the indel sites on genomic DNA. (**B**) Western blot analysis of control and Cecr2 knockout (sg1 and sg2) 4T1 cells, with the detected mouse CECR2 band at ~55 kDa. (**C-D**) Representative images (**C**) and quantification (**D**) of colony formation assays comparing control and Cecr2 knockout 4T1 cells. (**E**) Representative DAPI staining images from transwell invasion assays comparing control and Cecr2 knockout 4T1 cells. Scale bars: 200 μm. (**F**) Kaplan-Meier survival curve of immunodeficient BALB/c nude mice with tail vein injection of control (n=6) and Cecr2 knockout 4T1 cells (n=7) using bioluminescence signals at $1x10^8$ as the end point. Mantel-Cox log-rank test was performed to calculate the *p* value. (**G**) Western blot analysis of control 4T1, Cecr2 knockout 4T1, and Cecr2 knockout 4T1 with CECR2 reconstituted expression, with the detected mouse CECR2 band at ~55 kDa and human CECR2 bands at both ~170 kDa (full length) and ~55KDa (shorter isoform). (**H**) Kaplan-Meier survival curve of immunocompetent BALB/c nude mice with tail vein injection of control 4T1 (n=10), Cecr2 knockout 4T1 (n=10) and Cecr2 knockout 4T1 with CECR2 reconstituted expression (n=10) using bioluminescence signals of $1x10^8$ as the end point. Mantel-Cox log-rank test was performed to calculate the *p* values. (**I**) Tumors were scored based on the percentage of tumors in the lungs using the parameters described in **Figure S3E**. (**J**) Mammary tumor growth curve of immunocompetent BALB/c mice orthotopically injected with control (n=6) and Cecr2 knockout 4T1 cells (n=7). (**K**) Lung metastasis lesions from the mice described in (**J**) were quantitated using metastatic tumor scores with the parameter as **Figure S3E**. ** $p < 0.01$, *** $p < 0.001$, *n.s.*, not significant. Representative data from triplicate experiments are shown, and error bars represent SEM.

### Supplemental Figure 5

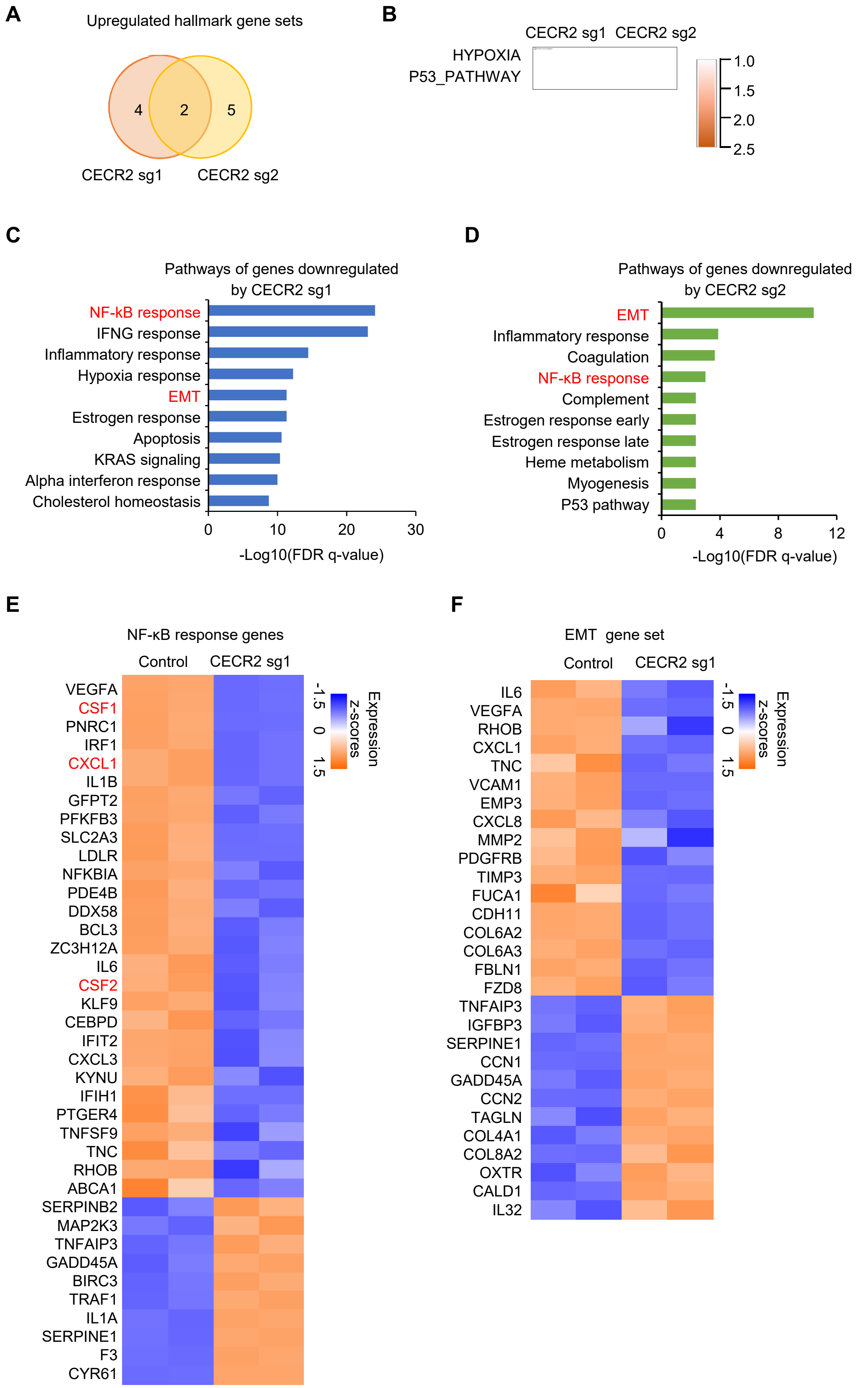

**Supplemental Figure 5. Pathways affected by CECR2 knockout in LM2 cells.**

(**A-B**) Gene set enrichment analysis comparing transcriptomes of CECR2 knockout (CECR2 sg1 and CECR2 sg2) with control LM2 cells. Venn diagram (**A**) showing the number of shared upregulated hallmark pathways (**B**). (**C-D**) Top 10 downregulated pathways in CECR2 knockout cell lines (sg1 and sg2) by GO analysis. (**E-F**) Heatmap of deregulated NF-κB response and EMT genes upon CECR2 depletion.

### Supplemental Figure 6

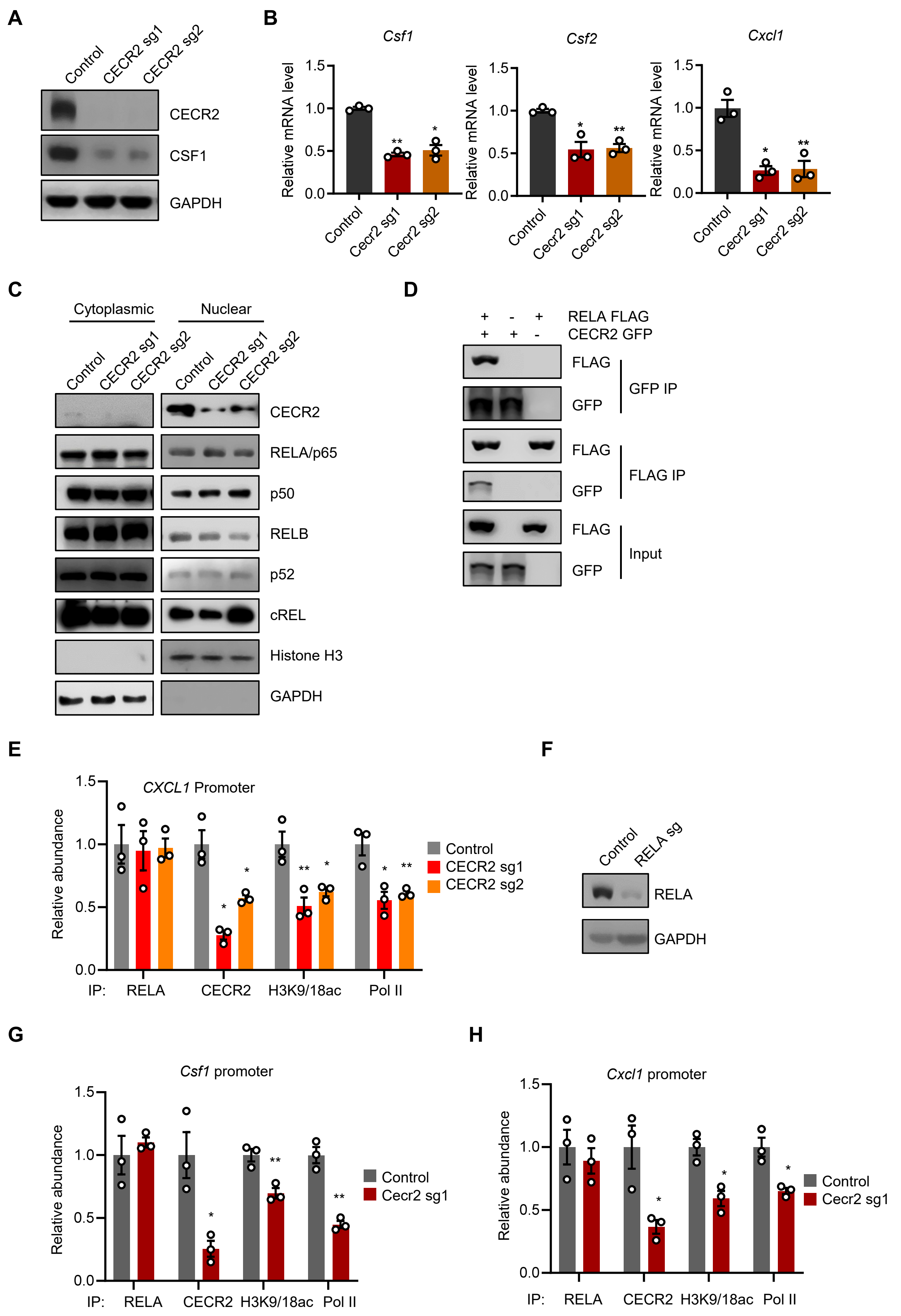

**Supplemental Figure 6. CECR2 interacts with RELA to activate NF-κB response genes.**

(**A**) Western blot analysis of control and CECR2 knockout (sg1 and sg2) LM2 cells. (**B**) RT-qPCR analysis of *Csf1, Csf2* and *Cxcl1* with control and cecr2 knockout (sg1 and sg2) 4T1 cells treated with 20 ng/ml TNF-α for 3 hours. (**C**) Western blot analysis of NF-κB family members in control and CECR2 knockout LM2 cells. (**D**) Western blot analysis of cell lysates (Input) and anti-immunoprecipitates (IP) from HEK293T cells transfected with the indicated plasmids. (**E**) ChIP-qPCR analyses with the indicated antibodies of the *CSF1* promoter in control and CECR2 knockout (sg1 and sg2) LM2 cells. (**F**) Western blot analysis of control and RELA knockout (sg) LM2 cells. (**G-H**) ChIP-qPCR analyses with the indicated antibodies of the *Csf1* (**F**) and *Cxcl1* (**G**) promoters in control and Cecr2 knockout (sg1) 4T1 cells. *$p < 0.05$, ** $p < 0.01$. Representative data from triplicate experiments are shown, and error bars represent SEM.

### Supplemental Figure 7

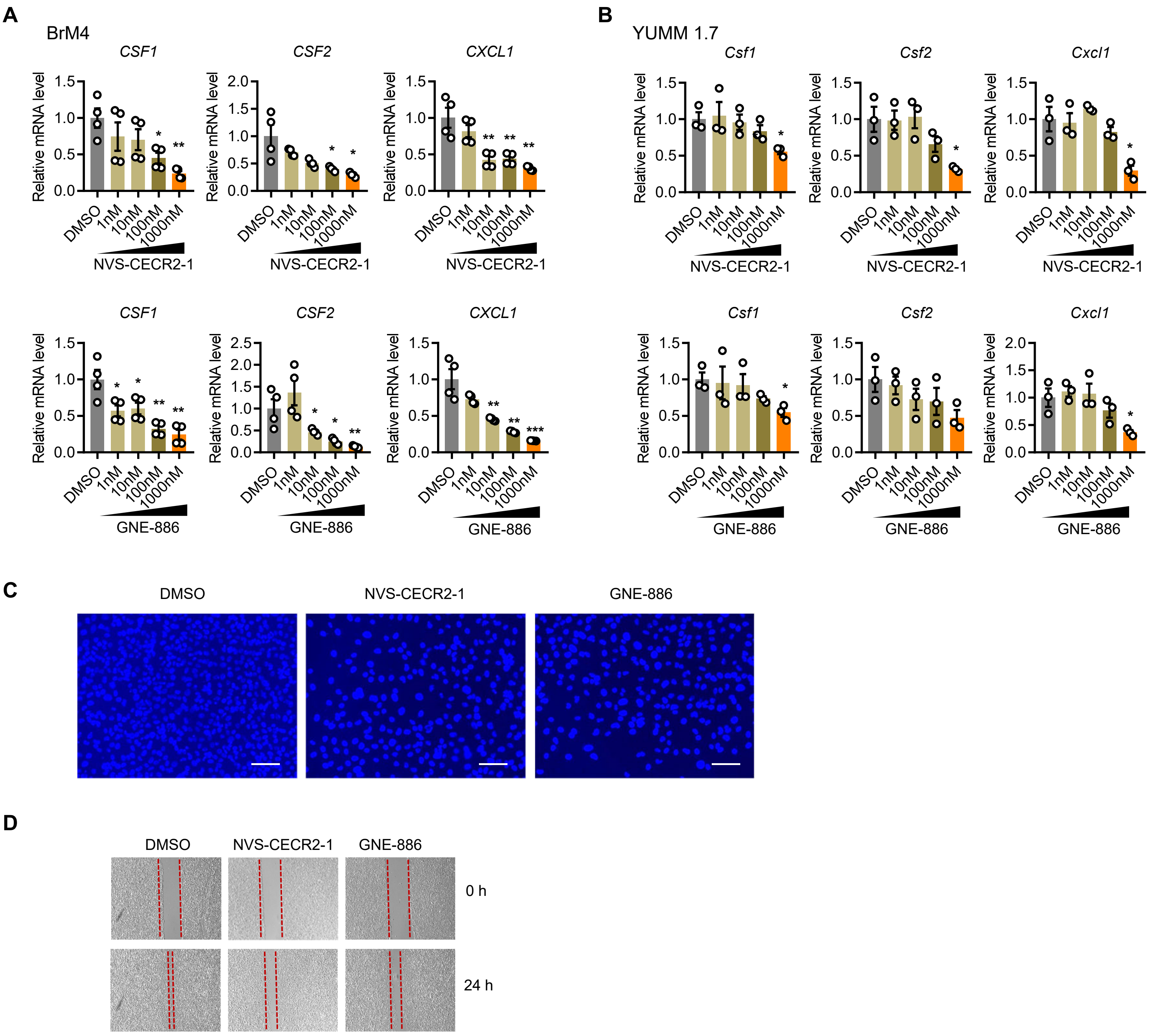


**Supplemental Figure 7. CECR2 inhibition suppresses the expression of NF-κB response genes as well as the migration and invasion capability.**
(**A**) RT-qPCR analysis of *CSF1, CSF2* and *CXCL1* expression in brain metastatic PC9-BrM4 (BrM4) lung cancer cells treated with DMSO, NVS-CECR2-1 and GNE-886 for 2 days and followed with treatment with 20 ng/ml TNF-α for 3 hours. * $p < 0.05$, ** $p < 0.01$. Representative data from triplicate experiments are shown, and error bars represent SEM. (**B**) RT-qPCR analysis of *Csf1, Csf2* and *Cxcl1* expression in YUMM1.7 metastatic melanoma cells treated with DMSO, NVS-CECR2-1 and GNE-886 for 2 days and followed with treatment with 20 ng/ml TNF-α for 3 hours. * $p < 0.05$, ** $p < 0.01$. Representative data from triplicate experiments are shown, and error bars represent SEM. (**C**) Representative DAPI staining images from transwell invasion assays comparing LM2 cells treated with DMSO, 1 μM NVS-CECR2-1 or 1 μM GNE-886 for 2 days. Scale bars: 200 μm. (**D**) Representative images showing wound healing distance at 24 hours in scratch assays comparing LM2 cells treated with DMSO, 1 μM NVS-CECR2-1 or 1 μM GNE-886 for 2 days.

#### Supplemental Figure 8

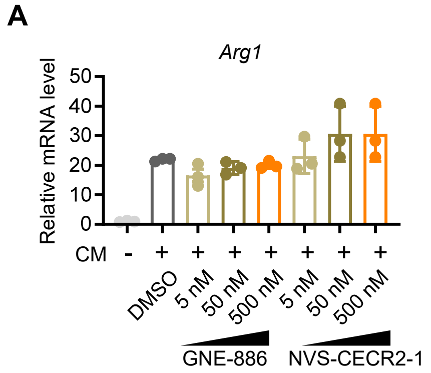


**Supplemental Figure 8. CECR2 inhibitors do not affect macrophage polarization *in vitro*.**

**(A-C).** Macrophages were seeded into 6-well plate and treated with conditioned media (CM) from 4T1 cells and DMSO, GNE-886 and NVS-CECR2-1 at indicated dosage for 2 days. RT-qPCR analyses of M2 macrophage markers *Arg1* **(A)**, *CD206* **(B)** *and IL-10* **(C)** were shown.

### Supplemental Figure 9

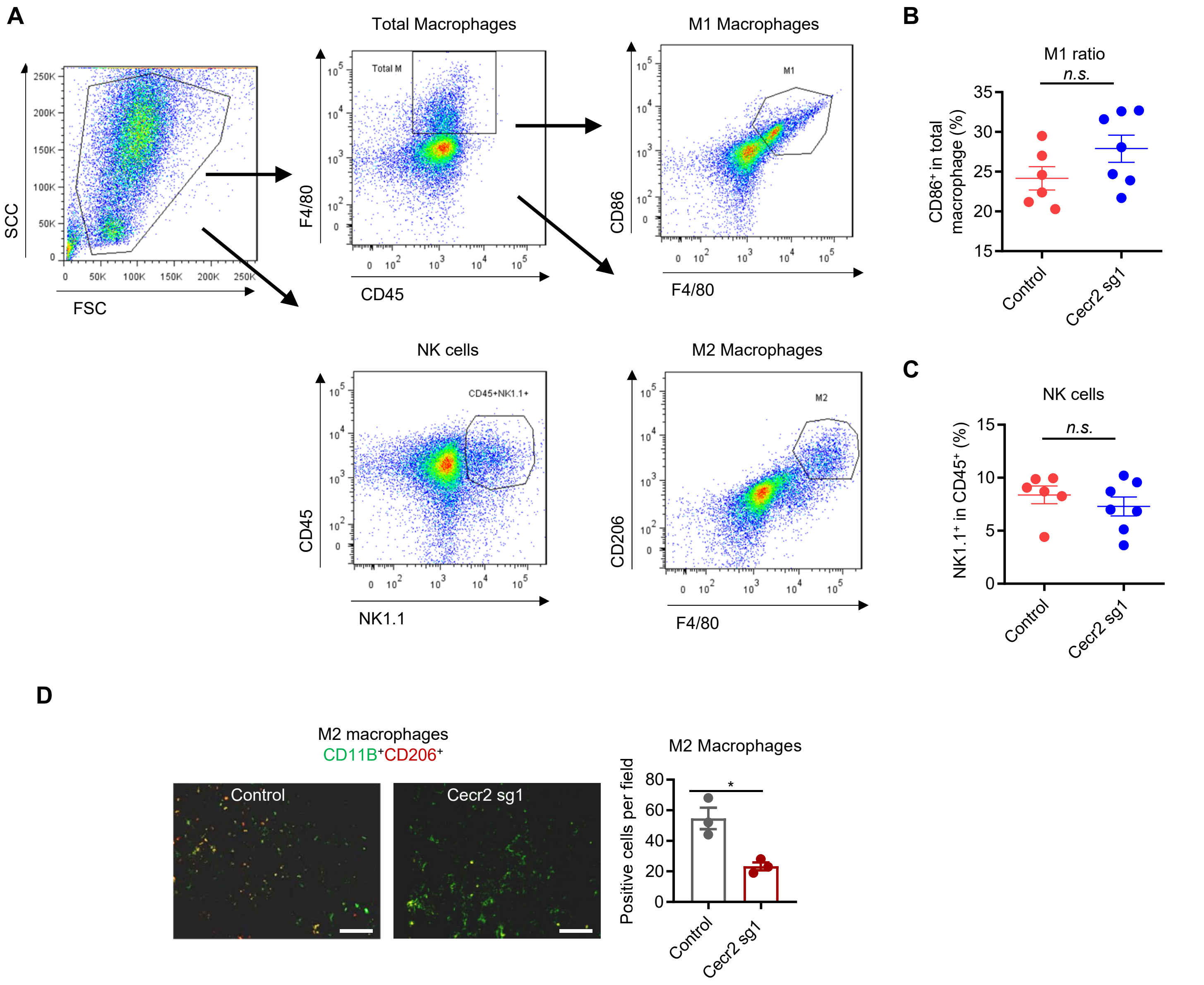


**Supplemental Figure 9. Cecr2 loss in 4T1 cells does not affect the percentages of M1-like macrophages and NK cells in lung metastases.**

(**A-C**) Flow cytometry analysis of TAMs and NK cells in the lungs from immunodeficient BALB/c nude mice with tail vein injection of control (n=6) and Cecr2 knockout (sg1) 4T1 cells (n=7) at week 2. Shown are representative flow cytometry plots (**A**), the ratios of M1 macrophages (**B**) and the percentages of NK cells (**C**). *n.s.* not significant. Representative data from triplicate experiments are shown, and error bars represent SEM. (**D**) IF staining of M2 TAMs in the lungs from immunocompetent BALB/c mice with tail vein injection of control (n=10) and Cecr2 knockout (sg1) 4T1 cells (n=10) at week 2. Shown are representative images (left panel) and quantification (right panel). Scale bars: 200 μm.

### Supplemental Figure 10

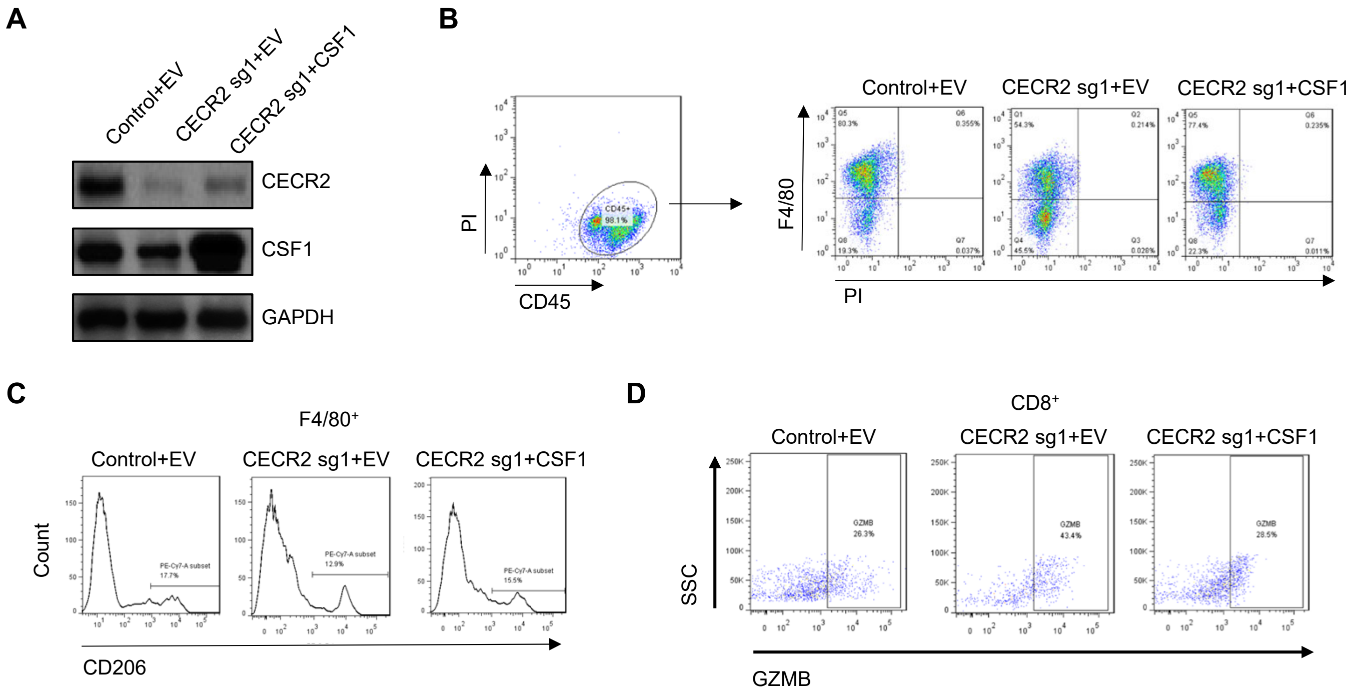


**Supplemental Figure 10. CECR2 activates CSF1 to inhibit anti-tumor immunity in metastatic microenvironment.**
(**A**) Western blot analysis of control 4T1, Cecr2 knockout (sg1) 4T1 cells, and Cecr2 knockout 4T1 cells with CSF1 overexpression. (**B-D**) Representative flow cytometry plots of total macrophages (**B**), M2 macrophages (**C**), and granzyme B expressing $CD8^+$ T cells (**D**) in the lung lesions of BALB/c mice injected with control 4T1, Cecr2 knockout (sg1) 4T1 cells, or Cecr2 knockout 4T1 cells with CSF1 overexpression. GZMB, granzyme B. Representative data from triplicate experiments are shown.
